# The 9p21.3 coronary artery disease risk locus drives vascular smooth muscle cells to an osteochondrogenic state

**DOI:** 10.1101/2024.05.25.595888

**Authors:** Elsa Salido, Carolina de Medeiros Vieira, José Verdezoto Mosquera, Rohan Zade, Parth Parikh, Shraddha Suryavanshi, Clint L. Miller, Valentina Lo Sardo

**Affiliations:** Department of Cell and Regenerative Biology; University of Wisconsin-Madison; Madison, WI 53705 USA; Department of Genome Sciences; Department of Biochemistry and Molecular Genetics; University of Virginia; Charlottesville, VA 22908 USA

## Abstract

**Background:** Genome-wide association studies have identified common genetic variants at ∼300 human genomic loci linked to coronary artery disease (CAD) susceptibility. Among these genomic regions, the most impactful is the 9p21.3 CAD risk locus, which spans a 60 kb gene desert and encompasses ∼80 SNPs in high linkage disequilibrium. Despite nearly two decades since its discovery, the role of the 9p21.3 locus in cells of the vasculature remains incompletely resolved.

**Methods:** We differentiated induced pluripotent stem cells (iPSCs) from risk and non-risk donors at 9p21.3 into vascular smooth muscle cells. We performed single-cell transcriptomic profiling, including co-embedding and comparison with publicly available human arterial datasets. We conducted functional characterization using migration and calcification assays and confirmed our findings on iPSC-VSMCs derived from additional donors. Finally, we used overexpression of *ANRIL* followed by gene expression analysis.

**Results:** We demonstrated that iPSC-VSMCs harboring the 9p21.3 risk haplotype preferentially adopt an osteochondrogenic state and show remarkable similarity to fibrochondrocytes from human artery tissue. The transcriptional profile and functional assessment of migration and calcification capacity across iPSC-VSMCs lines from multiple donors concordantly resemble an osteochondrogenic state. Importantly, we identified numerous transcription factors driving different VSMC state trajectories. Additionally, we prioritized *LIMCH1* and *CRABP1* as signature genes critical for defining the risk transcriptional program. Finally, overexpression of a short isoform of *ANRIL* in non-risk cells was sufficient to induce the osteochondrogenic transcriptional signature.

**Conclusions:** Our study provides new insights into the mechanism of the 9p21.3 risk locus and defines its previously undescribed role in driving a disease-prone transcriptional and functional state in VSMCs concordant with an osteochondrogenic-like state. Our data suggest that the 9p21.3 risk haplotype likely promotes arterial calcification, through altered expression of *ANRIL*, in a cell-type specific and cell-autonomous manner, providing insight into potential risk assessment and treatment for carriers.

**Highlights:** The 9p21.3 CAD risk locus promotes the transition of iPSC-derived Vascular Smooth Muscle Cells (iPSC-VSMCs) to an *osteochondrogenic phenotype*, both transcriptionally and functionally.

iPSC-VSMCs carrying the risk haplotype at 9p21.3 display a distinct transcriptomic signature, including osteochondrogenic markers *SOX9* and *COL2A1*, along with two novel markers: *LIMCH1* and *CRABP1*. Knockout of the entire haplotype reverts this signature to a non-risk state, demonstrating a *clear genotype-to-phenotype causal effect* of the risk allele.

Short isoform 12 of the lncRNA *ANRIL*, which partially overlaps the 9p21.3 locus, is sufficient to induce this transcriptional signature.

The iPSC-VSMC transcriptional profile strongly resembles that of *ex-vivo* human arterial VSMCs, providing a *human-specific model* to study VSMC phenotypic alterations and investigate the effects of large CAD haplotypes on vascular wall cells.

## Introduction

In the past two decades, GWASs have made significant advancements in identifying genetic variants associated with CAD^1–4^. However, translating this knowledge into functional and mechanistic insights remains a challenging task. One of the key bottlenecks is the lack of an appropriate model system to evaluate the causal link between human-specific genetic traits and the biology of the tissues and cell types implicated in the disease. As the majority of genetic variants associated with CAD reside in noncoding regions, they are predicted to impact gene regulatory mechanisms^5^. Furthermore, the lack of conservation across species makes it difficult to use standard animal models to dissect human-specific CAD risk loci^6^. How noncoding genetic risk factors for CAD influence the ability of vascular cells to acquire disease-prone states is not well defined.

The 9p21.3 CAD risk locus was discovered in 2007 through three independent CAD GWASs^7–9^. This 60 Kb genomic region is unique to primates and contains approximately 80 SNPs in strong linkage disequilibrium in most populations, presenting a challenge to pinpoint causal CAD risk variants^7–9^. The locus resides in a gene desert region and only partially overlaps with the extensively spliced long noncoding RNA (lncRNA) gene *ANRIL* (*CDKN2B-AS1*)^10^. The lack of conservation of this region across species has posed challenges for conducting *in vivo* studies; the 9p21.3 locus is poorly syntenic in rodents, and no study has identified a murine homolog of *ANRIL*^11^. A previous study reported the deletion of a putative syntenic region in the noncoding region downstream of the *Cdkn2a/b* genes on murine chromosome 4. These mice exhibited increased tumor incidence and increased aortic smooth muscle cell proliferation but not atherosclerosis^12^. Given the poor conservation of this genomic region across most mammals newer approaches that maintain the complexity of the human genome are urgently needed to study the function of the human 9p21.3 locus.

Vascular smooth muscle cells (VSMCs) are the most abundant cells in arteries and exhibit remarkable plasticity, characterized by their ability to transition to diverse cell states under physiological conditions and during CAD pathogenesis^13,14^. VSMCs are traditionally separated into two major cell states: 1) “contractile” quiescent cells that regulate vascular tone and 2) “synthetic” highly proliferative and highly migratory cells that produce extracellular matrix (ECM) and have reduced contractile markers expression^13–15^. Recent studies using single-cell RNA and ATAC (Assay for Transposase-Accessible Chromatin) sequencing of healthy and atherosclerotic coronary arteries have revealed more granular VSMC phenotypic states^16–21^. For instance, VSMCs can transition to macrophage-like and foam-like cells^22^, fibromyocytes^19^, SEM (Stem cell, Endothelial cell, and Monocyte intermediary state)^16^, fibrochondrocytes/chondromyocytes^20,21,23,24^ and osteoblast-like cells^23^. However, to date, the impact of the 9p21.3 CAD locus on VSMC states and transitions has not been described.

To determine the role of the 9p21.3 risk haplotype in influencing cell state changes in VSMCs, we used a system that we have previously generated and validated, based on induced pluripotent stem cells (iPSCs), haplotype editing and vascular smooth muscle cell differentiation^25^. Using this system, we conducted comprehensive single-cell transcriptional profiling to identify VSMC states and potential state trajectories mediated by the risk haplotype at the 9p21.3 locus. Here we show for the first time that, in an isogenic background, the risk haplotype at 9p21.3 drives VSMCs to acquire an osteochondrogenic-like state, both transcriptionally and functionally. Risk iPSC-VSMCs maintain VSMC identity while showing increased expression of *SOX9* and *COL2A1*, significant reduction in migration capacity and increased calcification. Our pipeline revealed specific transcriptional regulators driving different VSMC state trajectories and prioritized two signature genes, accompanying *SOX9* and *COL2A1*, *LIMCH1* and *CRABP1*, defining the 9p21.3-risk-driven VSMC state. Overexpression of pathogenic *ANRIL* isoforms, we previously described, is sufficient to elicit this transcriptional signature, suggesting that *ANRIL* has a major role in this phenotype. Moreover, our analysis revealed that the risk haplotype at 9p21.3 alters the expression of genes at other CAD loci, suggesting that an interconnected CAD gene network may increase risk susceptibility in concert. Our study causally links the 9p21.3 CAD risk locus to an osteochondrogenic state of VSMCs, known to be pathogenic prone and defines a specific transcriptional signature to investigate further in subjects carrying the CAD risk haplotype at 9p21.3. Our study also provides a human model to study the impact of large non-coding haplotypes in affecting VSMCs state transition and promoting CAD pathogenesis.

## RESULTS

### iPSC-VSMCs are reflective of ex vivo human SMCs and reveal a distinct transcriptomic landscape caused by the CAD risk haplotype at 9p21.3

To explore the impact of the 9p21.3 CAD risk locus on cell state transition and VSMCs plasticity, we leveraged a system based on induced pluripotent stem cells (iPSCs), we previously generated and characterized^25^. Briefly, iPSC lines were derived from both risk and non-risk homozygous backgrounds at 9p21.3. TALEN (Transcription Activator-Like Effector Nucleases) -mediated genome editing was used to delete the 60 kb CAD region at 9p21.3^25^. These isogenic iPSC lines share identical genetic backgrounds and differ exclusively in the 9p21.3 CAD region, allowing strict and controlled evaluation of the impact of 9p21.3 in determining cellular phenotypes. We established a collection of lines composed of four genotypes: risk cells with an intact haplotype (RR), isogenic risk knockouts (RRKO), non-risk cells with an intact haplotype (NN), and non-risk knockouts (NNKO). To control for potential variability across iPSC lines, we used two clones per genotype for parallel vascular smooth muscle cell (VSMC) differentiation. A total of 8 lines (carrying the 4 genotypes) were differentiated to VSMCs over the course of 17 days as we previously described^25,26^. At D17, the iPSC-VSMCs were harvested, and a total of ∼27,000 cells were used for simultaneous 10X Chromium single-cell 3’ gene expression sequencing assay (**Figure 1A**). After QC and removal of low-quality cells (detailed in Methods), we visualized the data by uniform manifold approximation and projection (UMAP) embeddings to evaluate gene expression patterns across different genotypes.

**Figure 1.**
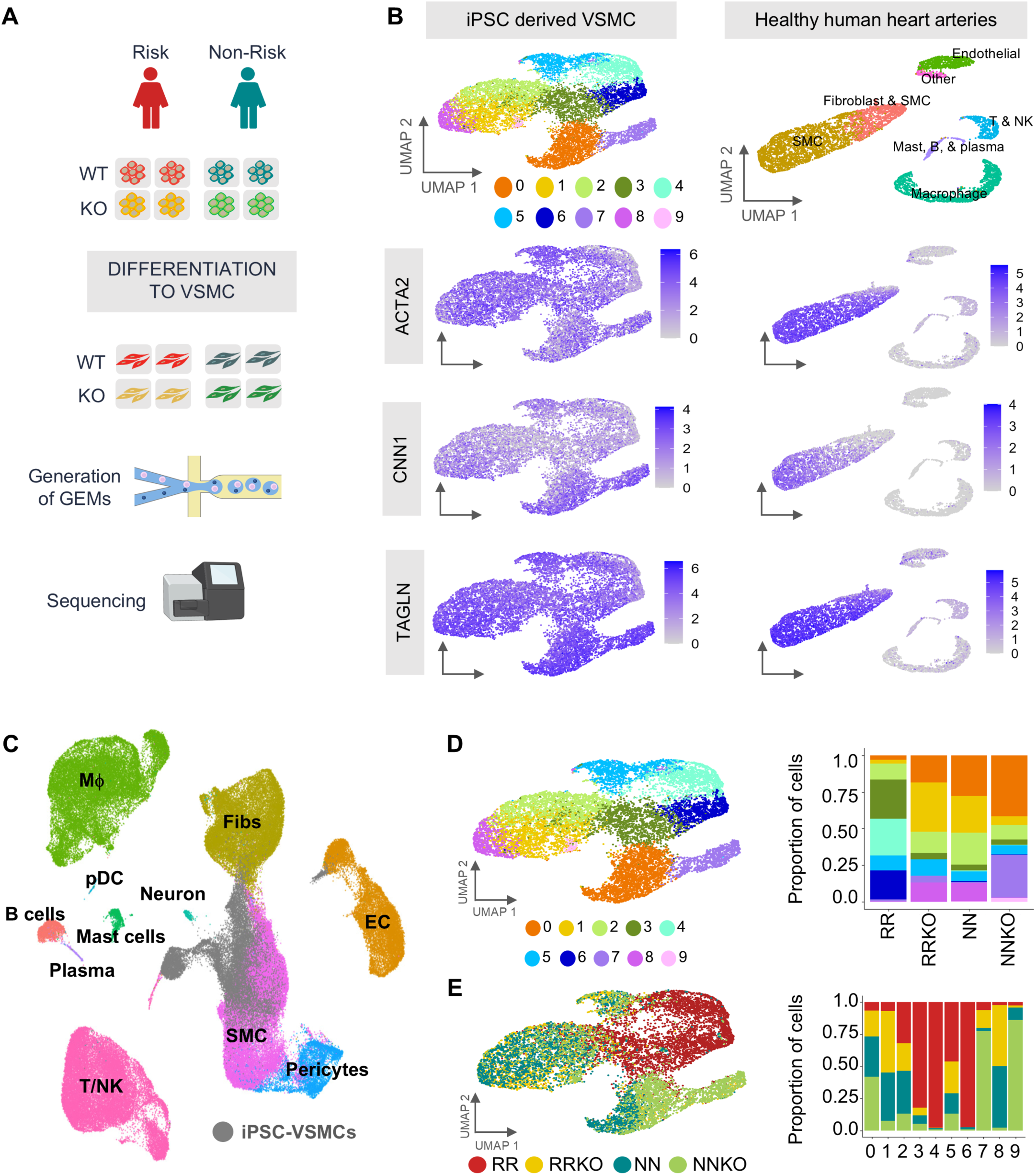
iPSC-VSMCs are reflective of *ex-vivo* human SMCs and reveal a 9p21.3 risk-specific transcriptional profile. **(A)** Schematic of experimental workflow. iPSC are from one homozygous risk and one homozygous non-risk donors, both with intact 9p21.3 locus. Isogenic knockout lines for each genotype were derived by TALEN-mediated haplotype editing to delete the entire CAD region. Two clones/genotype (N=8) were differentiated to VSMCs and used for single-cell sequencing. **(B)** UMAPs of scRNAseq data from iPSC-VSMCs collected at day 17 in 4 genotypes (RR-NN-RRKO-NNKO) (left) and UMAPs for *ex-vivo* healthy human artery data from Hu et al, 2021 (right). Topmost UMAPs are colored by unbiased clustering in the iPSC-VSMCs and by cell type in the *ex-vivo* data. Other UMAPs show expression of SMC markers in both datasets. **(C)** Projection of iPSC-VSMCs in the Metaplaq atlas of *ex vivo* human artery cells from Mosquera et al., 2023. **(D)** UMAP shows iPSC-VSMCs single cell transcriptome colored by unbiased clustering (left) and relative quantification of cluster contribution to each genotype (right). **(E)** UMAP shows iPSC-VSMCs transcriptome colored by genotype (left) and relative quantification of genotype contribution to each cluster (right).

To establish the robustness of this *in vitro* system in mimicking endogenous cellular phenotypes, we compared gene expression patterns with publicly available datasets of human heart arteries^17^ and evaluated the expression levels of established SMC markers, namely, *ACTA2*, *CNN1*, and *TAGLN* (**Figure 1B**). After *de novo* clustering and cell type annotation of the human artery data (**Figure S1A)**, we confirmed that iPSC-VSMCs represent an overall homogenous population expressing VSMC markers (**Figure 1B**). To further support the validity of iPSC-VSMC model, we leveraged the recently established ‘MetaPlaq’ atlas, a multidataset scRNA-seq atlas of human coronary and carotid artery cells^20^. Batch integration and co-embedding of the iPSC-VSMCs and the human artery scRNA-seq atlas in a low dimensional space showed strong qualitative overlap between iPSC-VSMCs and the *ex vivo* SMC cluster (**Figure 1C**). We detected minimal overlap only of a few cells with the fibroblast population (Fibs) and a small subset of iPSC-VSMCs projected on the endothelial cell (EC) cluster, indicating a remarkable similarity of the iPSC-VSMCs with endogenous human vascular SMCs.

We performed an unsupervised clustering analysis including all genotypes of iPSC-VSMCs and detected 10 transcriptionally distinct cellular clusters (**Figure 1D**). Although iPSC-VSMCs exhibit overall homogeneous expression of cell type-specific markers, this analysis showed that our system can detect different cell subpopulations, potentially representing different cellular states. To establish how different 9p21.3 haplotypes may influence the transcriptional landscape of VSMCs, we analyzed and quantified the cluster composition of our iPSC-VSMCs sorted by genotype (**Figure 1E**). Remarkably, cluster composition analysis revealed that clusters 0 and 1 account for ∼50% of the NN, NNKO, and RRKO samples, while the RR cells are mostly devoid of these clusters. Clusters 2 and 5 are equally represented across all genotypes. While NN, NNKO, and RRKO cells share similar cluster patterns, 72% of RR VSMCs comprise clusters 3, 4, and 6 (**Figure 1E**). RR cells tend to group together and away from other genotypes (NN, NNKO, RRKO), which lack the risk haplotype at 9p21.3 (**Figure 1E**). The remaining genotypes exhibit remarkable intermixing, suggesting that the deletion of the 9p21.3 CAD risk region (RRKO) mostly resembles a NN-like status. These data highlight the presence of transcriptionally independent cellular clusters in the overall population of iPSC-VSMCs, with three clusters (3, 4, 6) populated exclusively (∼80%) by VSMCs carrying the risk haplotype at 9p21.3, indicating potential distinct cellular states. Harmony correction across samples minimally impact cluster composition (**Figure S2G-I**). Analysis of the different cell lines showed comparable cluster contribution across independent clones and strong correlation across clones of the same genotype (**Figure S2A-E)** and equal cell differentiation and density at this differentiation stage (**Figure S2F**).

Since the 9p21.3 CAD region resides close to the cyclin-dependent kinase inhibitors *CDKN2A* and *CDKN2B* (**Figure S3A-B**), we evaluated the expression of these genes in the distinct clusters in VSMCs, which could possibly explain the divergent state. Although both genes showed variable expression in the different clusters, they did not localize to any specific genotype and were not enriched in RR VSMCs, excluding the possibility that the difference in the RR VSMC transcriptomic profile is due to genotype-specific expression of these genes (**Figure S3A-B**). To address the contribution of additional neighboring genes, we extended our analysis to a region spanning ∼ 2Mb upstream and ∼3 Mb downstream the CAD haplotype at 9p21.3. *KLHL9, FOCAD, FOCAD-AS1, LINC01239, DMRTA1, MTAP, HACD4, MLLT3* reached the level of detection, but did not reveal specific enrichment or considerable differences across cellular clusters and did not localize with the three unique RR VSMCs clusters (**Figure S3C**), suggesting that genes in the 9p21.3 region do not substantially contribute to the RR VSMC phenotype.

### The 9p21.3 risk haplotype reduces VSMC plasticity

To identify cluster-specific gene expression signatures, we conducted differential expression analysis (DEA) and annotated the top 10% cluster-specific genes (**Figure 2A, Table S1**). We highlighted a set of genes whose pattern of expression is unique in all risk clusters (**Figure 2A inlet, Table S2**). Gene Ontology (GO) enrichment analysis (**Table S3**) on all significant genes (Bonferroni corrected *p*-value < 0.05) driving each of the 10 clusters revealed two main patterns of biological processes in the NN, NNKO, and RRKO cells: (1) clusters 5 and 7 are enriched for cell cycle-related processes and concomitantly depleted of extracellular matrix organization and tissue remodelling processes; (2) clusters 0, 1, 2, and 8 are enriched in muscle development, wound healing and adhesion processes and depleted of cell cycle-related processes (**Figure S4**). Remarkably, this analysis revealed a different trend in RR VSMCs. Several terms related to healthy SMC function, such as cell adhesion and wound healing, were among the most significantly downregulated in two of the three risk clusters (4-6). This trend was accompanied by enriched cell cycle genes in cluster 4 and non-SMC lineages, including genes related to cartilage development and axonogenesis in clusters 3 and 6, respectively (**Figure 2B**). Interestingly, Cluster 6 exhibited strong enrichment for neuron projection-related processes.

**Figure 2.**
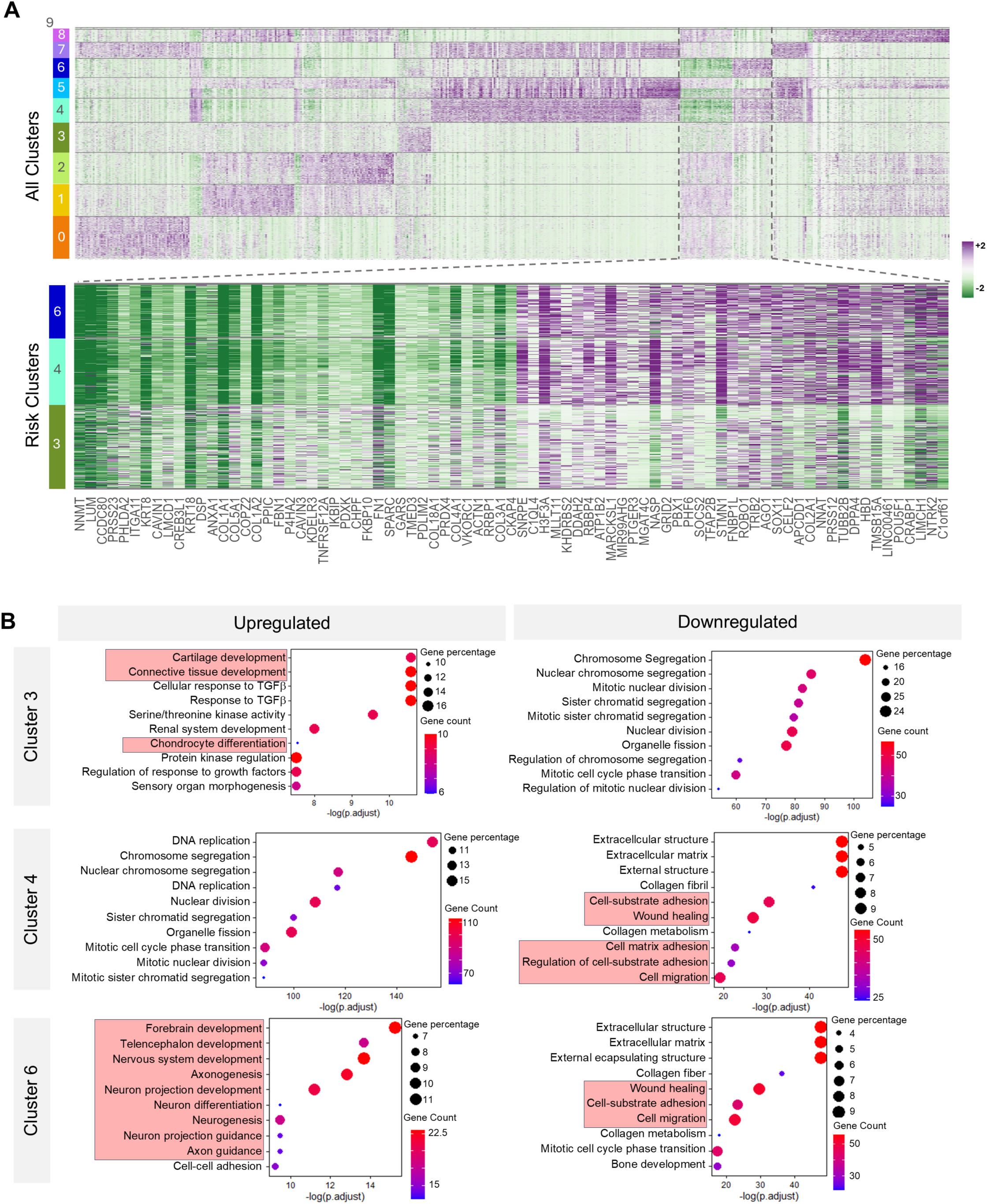
Differential gene expression analysis and Gene Ontology Enrichment reveal unique patterns of expression in RR VSMCs. **(A)** Top heatmap shows the top 10% genes by adjusted *p*-value driving each cluster (based on differential expression analysis) as they are expressed across all clusters. Bottom heatmap shows genes expressing uniformly and uniquely among risk clusters. **(B)** The top 10 gene ontology terms enriched in the differentially expressed genes driving each RR cluster (3-4-6). On the left GO terms enriched in the upregulated genes, on the right GO terms enriched in the downregulated genes are shown. Highlighted in red are pathways of interest.

To assess potential divergent cell differentiation in RR iPSCs we performed a closer examination of the axonogenesis and neuron projection genes using the GTEx human scRNA-seq atlas^27^. We analyzed the expression pattern of these genes across different tissues, including arteries and several tissues with high SMC content (Ovary, Bladder, Lung, etc.) (**Figure S5A**). We detected global expression of these genes across many tissues, suggesting that the enrichment in neuron-like GO terms is a result of biased functional annotation of most of these genes, rather than an alternative differentiation of RR VSMCs into neuron-like derivatives. Further, we did not observe significant enrichment of neuronal-specific genes in the risk cells (**Figure S5B, Table S4**). Thus, the presence of gene signatures related to axon guidance and neuronal projections may instead reflect morphological changes occurring in RR VSMCs, we have previously described^25,28^.

VSMCs are known to exhibit high plasticity and have been shown to acquire numerous transitional states^14,29,30^. Given the distinct gene expression signatures in RR VSMCs, we next investigated whether different genotypes at 9p21.3 could drive divergent transcriptional trajectories. We independently evaluated the dynamics of cell cluster transitions across each genotype using pseudotime analysis in Monocle3 (**Figure 3A**). We hypothesized that RR cells would follow a divergent cell trajectory, while the other three genotypes would converge on a similar trajectory. We selected cluster 2 as a common anchor for the trajectory analysis, given the balanced cell compositions by all four genotypes in this cluster and we evaluated the genotype specific trajectory of gene expression. The NNKO and RRKO lines converged on the same trajectory, predicting that the absence of the 9p21.3 haplotype has an equal effect on transcriptional states, independent of the genetic background. The *KO trajectory* is characterized by a transition from cluster 2 through 0 and terminating at 7. Gene ontology analysis of these clusters indicated a transition from a quiescent (clusters 2 and 0) to a more proliferative state (cluster 7) (**Figure S5C, Table S5**). The *nonrisk trajectory* mirrors the KO transition except for the last stage, terminating in cluster 0 instead of 7, suggesting that fewer NN cells transition to a proliferative (synthetic-like) state than NNKO or RRKO cells (**Figure 3A-B and Figure S5C**). Analysis of the *risk trajectory* substantiated our hypothesis of divergent state acquisition for RR VSMCs. The risk states populate the upper right portion of the UMAP, moving from cluster 2 through clusters 3, 4, and 6. Gene Ontology showed increasing non-VSMC signatures in RR VSMCs, suggesting that the risk haplotype drives VSMCs to acquire a divergent state (**Figure 3A-B**). To determine potential drivers of these trajectories, we interrogated the transcriptional trajectory to identify transcription factors potentially mediating these different cell states. We identified a “*common*” trajectory module including *MXD4, HOPX, PITX2, HAND2*, *HMGB2, MXD3, FOXP1*; a “*not risk*” module, shared by NN, NNKO, RRKO cells, including *CEBPD, HMGB3, TCF19, PSIP1, PHF19, MSX2*; and a “*knockout*” module including *SOX4, FOXM1*. Interestingly, we identified numerous transcription factors participating in the risk trajectory. The “*risk*” module comprises *BHLHE40, PBX1, SOX11, GTF21, CAMTA1, E2F2, POU3F1, ILF2, SOX2, IRX2, ID4, SP8* (**Figure 3C, Table S6**).

**Figure 3.**
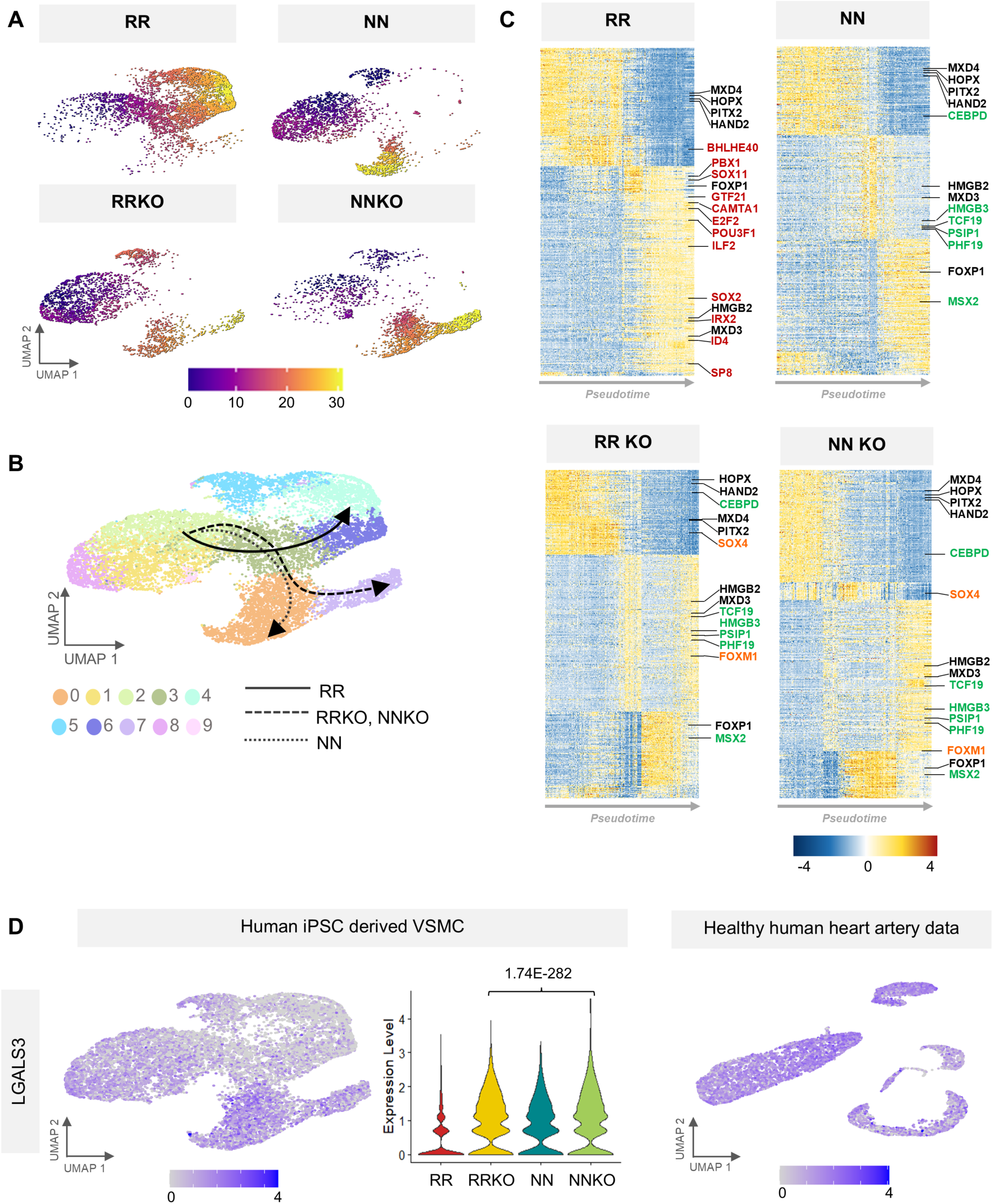
9p21.3 RR VSMCs show a divergent state trajectory. **(A)** Pseudotime analysis for each genotype. **(B)** Reconstruction of each genotype trajectory superimposed on a UMAP colored by cluster. **(C)** Heatmap showing the top 500 genes driving the trajectory of each genotype. Genes annotated are transcription factors (TFs). TFs in black are common to all trajectories, red TFs are only present in the risk trajectory, green TFs are present in all genotypes lacking the risk haplotype (NN, NNKO, RRKO), and orange TFs are only present in the knockout trajectories. **(D)** UMAP showing expression of *LGALS3* in iPSC-VSMCs (left), violin plot showing the expression of *LGALS3* across genotypes with *p*-value calculated via non-parametric Wilcoxon rank sum test (middle), and UMAP of *LGALS3* expression in healthy human artery data from Hu et al., 2021 (right).

Lineage tracing studies in mice and, more recently, single-cell transcriptomic analysis of human artery samples, have described numerous VSMC state nuances^14,29,30^. To investigate whether the cell state transition of our *in vitro* system recapitulates previously identified VSMC states or phenotypic switches, we analyzed the expression of known master regulators of VSMC phenotypic modulation. We sought to analyze traditional modulators of VSMC phenotypic switch, MYOCD and SRF^31,32^. MYOCD was detected only in RR and RRKO VSMCs, while SRF was expressed relatively evenly across genotype. Moreover, the stem cell marker KLF4^33–35^ was not detected in our dataset (**Figure S6A**). Overall, the expression patterns of these transcriptional regulators, which were previously identified to play a role in VSMC modulation, do not differ significantly by genotype or SMC cluster, suggesting that the risk phenotype may be driven by a different transcriptional program. We then interrogate our dataset for evidence of transdifferentiation to other cell types or aberrant CAD-related cell states^14,16,19,20,29,35^. We used a single-cell dataset from healthy human arteries^17^ together with two additional published datasets using endogenous atherosclerotic human arteries^16,19^ (**Figure S1B-C)** and looked for evidence of either transition to fibromyocytes and SEM (Stem cell, Endothelial cell, and Monocyte intermediary state), or transdifferentiation to other cell types, including macrophages or endothelial cells. We did not detect expression of known fibromyocyte markers (*TNFRSF11B or VCAM1*) (**Figure S6B**) in iPSC-VSMCs overall, precluding the possibility that RR VSMCs acquire one of these known VSMC states. Similarly, we observed a lack of expression of macrophage (*CD68* and *AIF1*) or endothelial cell (*PECAM1* and *vWF*) (**Figure S6C**) markers, suggesting that the RR VSMCs are unlikely to originate from transdifferentiation to other cell types. Overall, this analysis shows no significant resemblance between the RR cells and some of the previously described cell types and states we surveyed.

In mouse models of atherosclerosis, *Lgals3* was previously linked to macrophage-like VSMCs during atherosclerosis progression^36^. More recently, *Lgals3* has been described as a marker of VSMC plasticity and an indicator of a transitional state for subpopulations of VSMCs that undergo phenotypic modulation^34^. Interestingly, *LGALS3* was uniformly expressed in all clusters populated by RRKO, NN and NNKO cells, while the expression of *LGALS3* was significantly decreased in RR cells (**Figure 3D**). This pattern suggests that NN, NNKO and RRKO cells exhibit a dynamic transitional phenotype, while RR cells likely lack plasticity and do not undergo known phenotypic modulation.

To assess whether other genes within known CAD genomic loci are affected in the iPSC-VSMC RR population, we analyzed the expression of 90 genes residing in known CAD loci^37^. We found 19 genes significantly differentially expressed between the RR and other genotypes that were homogeneously expressed or repressed in RR VSMCs (**Figure S7, Table S7**). Individual cluster survey of these genes shows a consistent trend of upregulated genes in all three risk clusters (3,4,6) and homogeneous downregulation of most of the genes, except for *TNS1, PROCR, MRAS and FURIN*, showing slightly different level of expression across the three clusters. Overall, this analysis suggests that the 9p21.3 locus influences multiple other genes linked to CAD and that the risk status of the cells may arise from a cumulative effect of gene expression alterations at other genomic loci.

### Risk iPSC-VSMCs exhibit an osteochondrogenic-like state characterized by novel CAD signature genes

The GO analysis indicated an enrichment in connective and cartilage lineages, hence, we sought to explore the possibility that RR VSMCs acquire an osteo/chondrogenic state. *SOX9* and *RUNX2* are master regulators of bone and cartilage differentiation and have been shown to be upregulated, in CAD vessels, in VSMCs transitioning toward osteogenic- and chondrogenic-like cells, promoting calification^23,24,29^. Recently, *SOX9* has been linked to vascular stiffening and aging through its modulation of collagen expression^38^. Evaluation of key markers for osteo- and chondrogenic lineages between RR cells and their isogenic RRKO cells revealed overall increased expression of both lineage markers in the presence of 9p21.3 risk (**Figure 4A**). Remarkably, *SOX9* was uniformly expressed at higher levels in RR cells compared to RRKO cells (**Figure 4B-C**). Additionally, *SOX9* upregulation was accompanied by significant upregulation of *COL2A1* and concomitant downregulation of *COL1A1* and *COL1A2*, suggesting that the cells acquire a chondrogenic fate and alter ECM production (**Figure 4B-C**).

**Figure 4.**
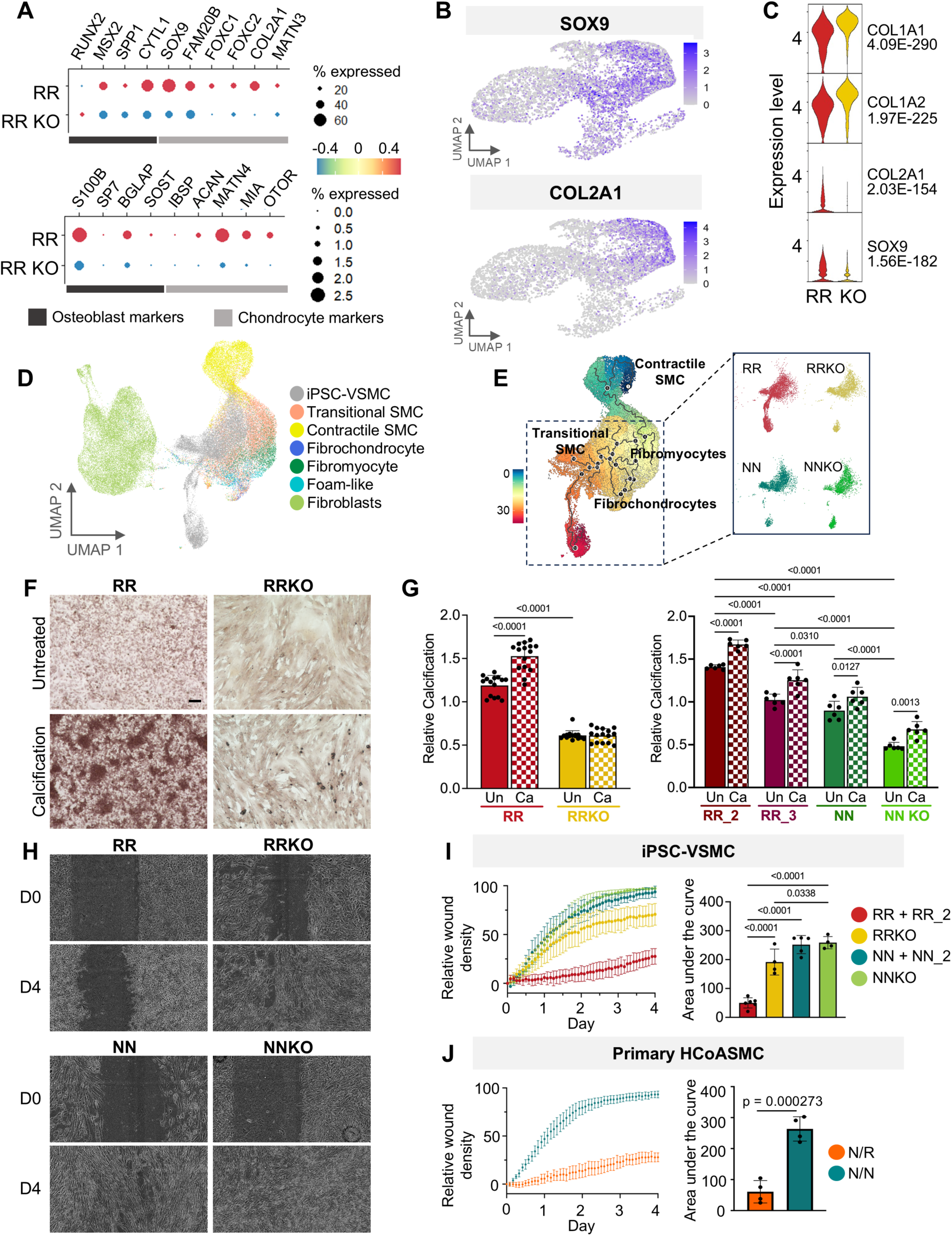
RR VSMCs acquire an osteochondrogenic-like state. **(A)** Bubble plots of osteoblast and chondrocyte markers in RR and RR KO. **(B)** UMAPs showing upregulation of *SOX9* and *COL2A1* in risk VSMCs. **(C)** Violin plots showing expression of *SOX9*, *COL2A1*, *COL1A1*, and *COL1A2* in RR VSMCs and isogenic RRKO, with *p*-values calculated by non-parametric Wilcoxon rank sum test. **(D)** Secondary annotation of iPSC-VSMCs and *ex-vivo* SMC/Fibroblasts (from Mosquera et al., 2023) harmonization showing modulated subpopulations. **(E)** Pseudotime analysis of harmonized iPSC-VSMCs and Metaplaq SMC/Fibroblasts showing the trajectory of iPSC-VSMCs and profiles of specific genotypes in the insert to the right. **(F)** Alizarin red staining of iPSC-VSMCs from RR and RRKO genotypes. Cells were untreated or treated for 7 days with a calcification media. Scale bar is 100μm. **(G)** Quantification of Alizarin red for RR cells and RRKO (left); N=15 (5 fields of view for three independent calcification experiments) (left). Alizarin staining for risk VSMCs from two additional donors (RR_2, RR_3), NN and NNKO lines (right); N=6 (3 fields of view for two independent calcification experiments). Data are presented as mean±SD. *P*-value is calculated with One-way ANOVA with Bonferroni correction. **(H)** Wound/scratch assay of iPSC-VSMCs with different genotypes at 9p21.3 over time. **(I)** Graph of cell migration assay for iPSC-VSMCs of all genotypes (N=4 for RRKO and NNKO and N=6 for RR and NN including the isogenic RR and NN lines and 2 additional donors; RR_2 and NN_2). Breakout of the graph in Figure S9. Wound resolution is plotted over time (left) and statistical analysis of area under the curve (AUC) is on the right. Data are presented as mean±SD and *p*-value is calculated via One-way ANOVA with Bonferroni correction. **(J)** Cell migration assay for Primary HCoASMCs plotted over time. Data are from 4 cell lines derived from 4 independent donors, two per genotype (Heterozygous risk/non risk= NR and homozygous non risk=NN). AUC is on the right. Data are presented as mean±SD and *p*-value is calculated via unpaired *t*-test.

Using the most comprehensive human artery single-cell atlas available, MetaPlaq^20^, we examined whether iPSC-VSMCs resemble VSMC states in *ex vivo* arterial samples. Integration of SMCs and fibroblasts from the atlas with iPSC-VSMCs revealed a remarkable overlap of iPSC-VSMCs with *ex vivo* arterial SMCs (**Figure S8A-B**). Using fine-grained cell annotations in this atlas, we demonstrated that iPSC-VSMCs mostly overlap with Transitional SMCs, Fibromyocytes, Foam-like SMCs and Fibrochondrocytes (**Figure 4D**). Pseudotime trajectory analysis incorporating SMCs from human arteries and iPSC-VSMCs from this study showed that the non-risk iPSC-VSMCs (NN, NNKO, RRKO) mostly overlap with transitional populations of SMCs, corroborating the findings obtained using *LGALS3* as a marker of phenotypic modulation. Interestingly, RR cells resemble fibrochondrogenic SMCs and partially foam-like cells. Additionally, we identified a divergent cell population, mostly derived from RR VSMCs (**Figure 4E**), which may represent a less abundant transitional cell population in the current atlas.

Previous studies have shown that VSMC-derived osteochondrogenic cells promote calcification^21,23^. Alizarin red staining was performed on fully differentiated VSMCs with different genotypes at 9p21.3. In basal conditions, without any calcifying challenge, RR VSMCs show a ∼2-fold increase in calcium deposits, compared to the KO isogenic lines (RR Un=1.189+/-0.12 vs RRKO Un=0.612+/-0.05) (**Figure 4F-G**), corroborating the evidence presented thus far of an osteochondrogenic fate for RR VSMCs. Upon exposure to calcification conditions for 7 days^39^, RR VSMCs show a significant increase in calcification (RR Ca=1.527+/-0.16), while RRKO cells are unchanged (RRKO Ca=0.612+/-0.07). Importantly, we differentiated iPSCs from three additional independent donors^40^, including 2 risk iPSC lines (RR_2, RR_3) and 2 non-risk isogenic lines NN and NNKO. Our data showed consistent results with increased Alizarin staining in RR lines in basal condition compared to NN and NNKO lines, although showing different extent of calcification across donors (RR_2 Un=1.408+/-0.02; RR_3 Un=1.048+/-0.06; NN Un=0.897+/-0.11; NNKO Un=0.482+/-0.04). Upon the same calcification culture condition as previously showed, all lines show increased calcification (RR_2 Ca=1.671+/-0.05; RR_3 Ca=1.298+/-0.05; NN Ca=1.061+/-0.11; NNKO Ca=0.678+/-0.09) (**Figure 4G and Figure S9A**). These data corroborate a causal effect of the 9p21.3 risk haplotype in eliciting an osteochondrogenic state and show that the transcriptional profile of RR VSMCs is accompanied by functional evidence of pronounced calcification. Remarkably, this phenotype is consistent across iPSC-VSMCs derived from several human donors, demonstrating the robustness of this phenotype.

Migration capacity is an essential feature of VSMCs^14,41^ and has a dual role in CAD initiation and progression^14,35,42,43^. Synthetic VSMCs migratory potential contributes to the invasion of the arterial intima during the early stages of CAD. However, migration of VSMCs has also a protective effect in stabilizing atherosclerotic plaques and preventing rupture, a leading cause of myocardial infarction. Previous studies have suggested lower migration capacity in chondromyocytes^21^, a state of VSMCs promoting calcification, as well as in VSMCs with reduced expression of *Lgals3*^34^. To assess the migratory capacity of iPSC-VSMCs carrying different genotypes at 9p21.3, we employed a wound/scratch assay followed by live imaging to assess cell migration of VSMCs within a matrix-rich niche. RR VSMCs show significant defects in migration, while RR KO, NN, and NN KO cells exhibited comparable migratory potential (**Figure 4H-I and Figure S9B-C**). After 4 days NN, NNKO and RRKO reach approximately 70-97% resolution of the wound (NN= 93.7% +/- 5.9; NNKO= 97.2% +/- 1.08; RRKO= 70.7% +/- 11.1). RR VSMCs, however, show a significant reduction in wound resolution of 27.6% +/- 8.02. We differentiated a new set of iPSCs derived from two additional donors carrying the risk and non-risk haplotype^40^ at 9p21.3 (RR_2 and NN_2)and performed a migration assessment confirming our findings, demonstrating the robustness of this phenotype (**Figure 4I and Figure S9C-D**). To further validate this genotype-specific effect on native VSMCs, we performed the same assays on 9p21.3 genotype-matched human coronary artery SMCs (HCoASMC). By probing two 9p21.3 heterozygous risk (NR) and two homozygous non-risk (NN) cell lines, we found a significantly reduced migration capacity in cells carrying the risk haplotype at 9p21.3 (**Figure 4J**). Our results provide the first evidence that RR iPSC-VSMCs, from multiple donors, are impaired in migratory potential, concordant with a phenotypic transition into osteochondrogenic-like cells.

To better define the transitional state of the RR VSMC population and identify new signature genes of 9p21.3 risk-specific phenotypes, we conducted a differential expression analysis between RR cells and all other genotypes and selected the most differentially expressed genes in the risk cells by *p*-value and fold change. Closer observation of the top 50 DEGs revealed only 11 that were uniformly and uniquely expressed in the entire RR cell population. We prioritized two genes with the highest fold enrichment and *p*-value for expression in RR cells: *LIMCH1* and *CRABP1* (**Figure 5A**).

**Figure 5.**
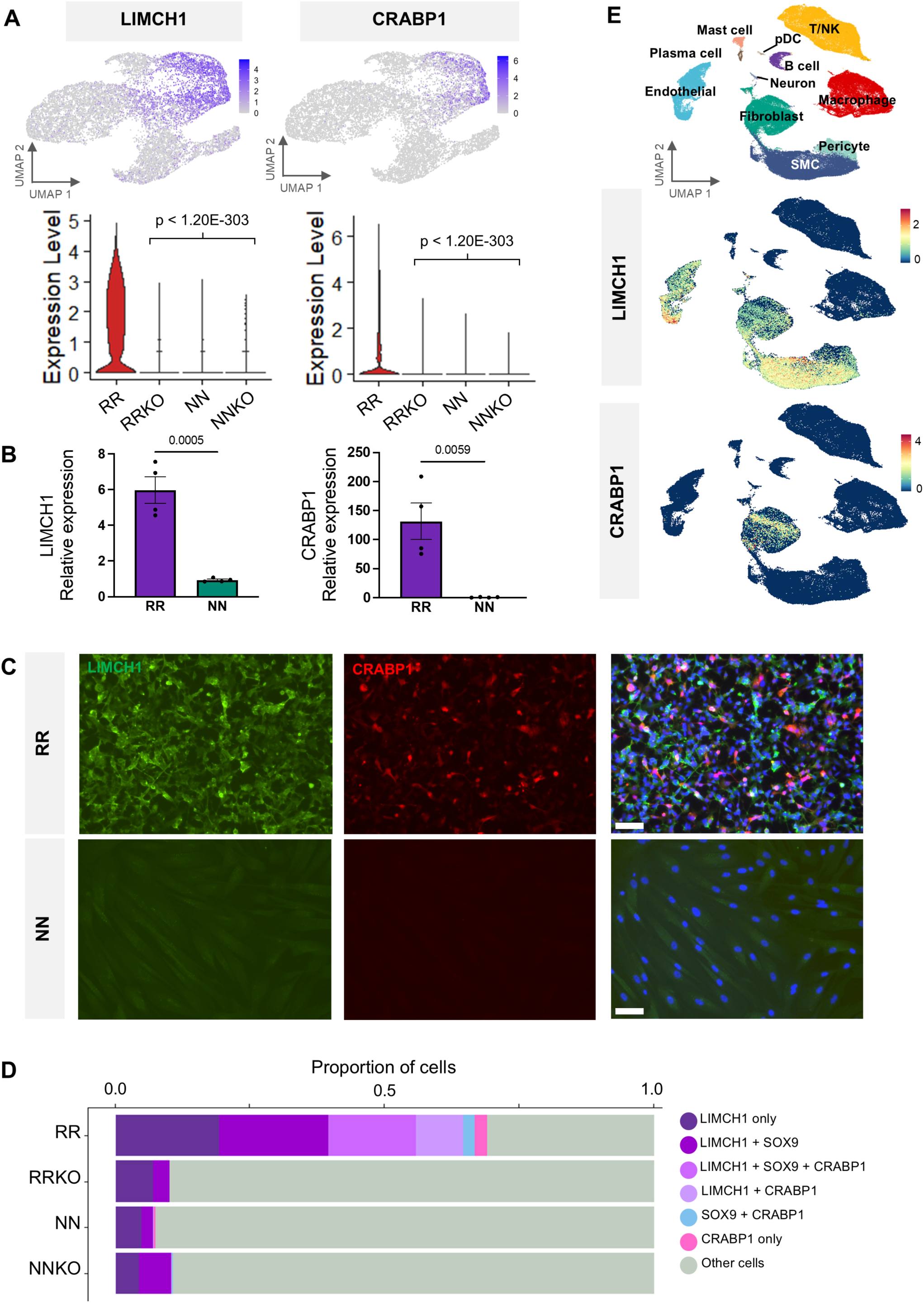
Identification of 9p21.3 risk VSMCs signature. **(A)** UMAPs showing upregulation of *LIMCH1* and *CRABP1* in iPSC-VSMCs (top), violin plots showing expression across genotypes (bottom). *p*-value calculated via non-parametric Wilcoxon rank sum test. **(B)** qPCR of *LIMCH1* and *CRABP1* enrichment in four additional donors (RR_2, RR_3 and NN_2, NN_3) in two independent VSMC differentiation experiments. N=4. Data are presented as mean±SD and *p*-value is calculated via unpaired *t*-test. **(C)** Immunocytochemistry staining for LIMCH1 and CRABP1 in RR and NN VSMCs. White bar indicates 50 μm. **(D)** Bar chart showing proportions of cells in different genotypes expressing *LIMCH1, SOX9*, and *CRABP1*. **(E)** Metaplaq atlas of human artery samples labeled with cell types (top), *LIMCH1* and *CRABP1* expression in Metaplaq atlas.

Immunocytochemistry confirmed that the expression of transcripts resulted in a high abundance of the LIMCH1 protein across the entire RR population and highlighted the expression of CRABP1 in ∼20% of RR VSMCs. The expression of these genes was not detected in NN cells, as predicted by the scRNAseq analysis (**Figure 5C**). To confirm the unique expression of these signature genes across multiple samples, we generated iPSC-VSMCs from two additional RR donors (RR_2, RR_3) and two NN donors (NN_2, NN_3)^40^. Gene expression analysis by qPCR confirmed that the expression of *LIMCH1* and *CRABP1* was greater in the VSMCs harboring the risk haplotype than in those harboring the NN haplotype (**Figure 5B**). Next, we analyzed the co-expression pattern of *LIMCH1* and *CRABP1* across the different genotypes. *LIMCH1*^+^ cells accounted for 64.7% of the RR VSMC population. Analysis of co-expression with the other signature genes revealed that 19.2% of RR VSMCs are exclusively expressing *LIMCH1*, 20.3% are *LIMCH1*^+^-*SOX9*^+^, 16.4% *LIMCH1*^+^-*SOX9*^+^-*CRABP1*^+^ and 8.8% are *LIMCH1*^+^-*CRABP1*^+^. RRKO, NN, and NNKO cells have 7-10% of cells expressing negligible levels of *LIMCH1* alone or *LIMCH1*+-*SOX9*+, indicating that this signature predominates with RR VSMCs (**Figure 5D**). Finally, we evaluated the expression of these genes in the MetaPlaq scRNA-seq atlas^20^ and found that *LIMCH1* and *CRABP1* were enriched in a few cell populations. *LIMCH1* can be found throughout the SMC population, with slight enrichment in the fibromyocyte population and a subpopulation of endothelial cells previously predicted to undergo EndoMT (**Figure 5E**)^20^. Interestingly, *CRABP1* was mostly detected in fibroblasts but was found in a subpopulation of cells predicted to resemble myofibroblasts^20^. Together, our analyses reveal signature genes in iPSC-VSMCs resembling novel unidentified cellular states in human arterial tissue, which warrant further investigation to assess their role in atherosclerosis.

### Isoform-specific ANRIL overexpression induces osteochondrogenic signature in VSMCs

Previously, we reported that *ANRIL*, the lncRNA that partially overlaps the CAD region at 9p21.3, has genotype-specific isoforms expression in iPSC-VSMCs harboring different genotypes at 9p21.3. Specifically, total *ANRIL* transcripts levels are higher in RR VSMCs compared to NN VSMCs. While long *ANRIL* isoforms, which terminate in exon 20, are equally expressed in RR and NN VSMCs, short *ANRIL* isoforms (terminating in exon 13) are expressed exclusively in RR VSMCs^25,28^. Here, we confirmed that the increased *ANRIL* expression is attributable to short *ANRIL* isoforms, as the long isoforms did not differ by genotype (**Figure S10A**). Concordant with our previous studies, differential *ANRIL* isoforms expression is robustly recapitulated in iPSC-VSMCs from additional RR and NN human donors (**Figure S10B**). To assess whether *ANRIL* has a causal effect in driving the osteochondrogenic state, we used doxycycline-inducible lentiviral particles carrying *ANRIL isoform 12*, a short ANRIL isoform, to exogenously induce *ANRIL* expression in fully 9p21.3 knockout cells (**Figure 6A**). NNKO cells were infected at D7 of the VSMCs differentiation with lentiviral particles carrying rTta (reverse tetracycline transactivator) and lentiviral particles carrying *ANRIL 12*. Cells not infected and treated with doxycycline and cells infected with rTta only were used as control. Doxycycline was initiated the day after and maintained until D17, when cells were harvested for gene expression analysis. Remarkably, upon overexpression of *ANRIL 12* in complete KO cells we observed significant upregulation of *SOX9, COL2A1, LIMCH1* and an increased trend for *CRABP1*, though not significant (**Figure 6A**). These results indicate that the expression of the *ANRIL 12 isoform* is sufficient to activate the expression of the 9p21.3 risk signature in NNKO cells.

**Figure 6.**
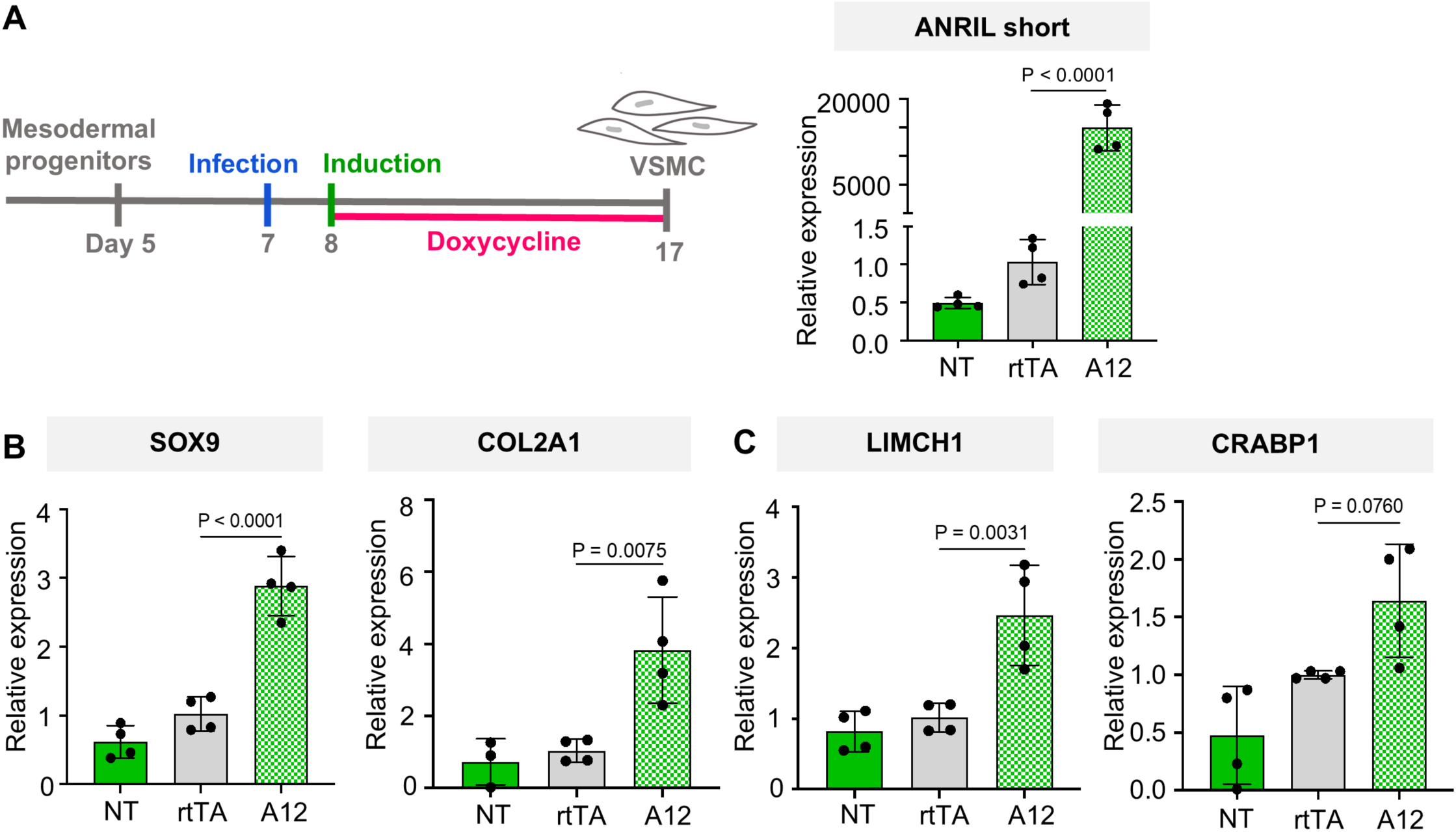
ANRIL induces osteochondrogenic signature genes. **(A)** Experimental diagram showing the *ANRIL isoform 12* overexpression experiment (left). Quantification via qPCR of *ANRIL* overexpression in NNKO cells not infected (NT), infected with rtTA (rtTA) and infected with rtTA+*ANRIL isoform 12* (A12) (right). **(B)** Quantification of the expression of key osteochondrogenic markers *SOX9* and *COL2A1* in response to *ANRIL 12* overexpression. Cells not infected (NT) and infected with only rTtA are shown as controls. **(C)** Quantification of the expression of risk markers *LIMCH1* and *CRABP1*. N=4. Data are presented as mean±SD and *p*-value is calculated by One-way ANOVA with Bonferroni correction.

## DISCUSSION

Genome-wide association studies (GWASs) have identified ∼300 genomic loci linked to CAD susceptibility^37,44,45^. However, the functions of these genomic regions, target genes, and relevant cell types that contribute to disease pathogenesis remain to be fully determined. Recent efforts using single-cell omics have identified CAD-associated cellular states in human coronary and carotid arteries^16,18–20^. However, how CAD-associated genetic variants and haplotypes impact cell state maintenance dynamics and cell plasticity in response to pathophysiologic stimuli is also unclear. Here, we present compelling evidence that the 9p21.3 CAD risk locus drives vascular smooth muscle cells (VSMCs) to adopt a state resembling osteochondrogenic cells. We identified a specific transcriptomic signature associated with 9p21.3 and evaluated this cell-state transition at both transcriptional and functional levels in iPSC-VSMCs from several risk and non-risk donors. Our findings demonstrate that 9p21.3 risk VSMCs exhibit a transcriptomic profile similar to fibrochondrocytes found in atherosclerotic human arteries. Consistent with this altered state, these cells show reduced migratory capacity and an increased propensity for calcification. Our study suggests that the 9p21.3 risk haplotype contributes to disease risk by inducing VSMCs to transition into a calcification-prone pathological state. This shift is mostly induced by *ANRIL*.

The 9p21.3 locus was the first CAD risk locus identified by GWAS almost two decades ago^7,8,46^, but its function remains to be fully defined. The risk haplotype frequency (∼50% of the population of non-African descent) and its strong additive effect for each risk allele account for ∼10-15% of the incidence of CAD in the US^47^, affecting ∼3 million people. Numerous studies have explored the function of the 9p21.3 locus using various approaches, including primary cultures^48–52^, cell lines^51,53^ and primary tissues^54,55^. However, new discoveries have been limited by a few important factors: i) they have mostly described correlations between the risk haplotype and cellular phenotypes without robust evidence of causality^56,57^; ii) most studies using primary cultures have predominantly employed cell types not primarily associated with atherosclerosis (whole blood, T cells, PMBCs)^48–51;^ and iii) functional studies on the 9p21.3 CAD locus employing genetic manipulation have been based on immortalized cell lines (HEK cells)^51^ or mouse models that lack synteny with the human 9p21.3 region^12^. In recent years, our group has provided evidence that the 9p21.3 CAD risk haplotype affects vascular smooth muscle cells (VSMCs) both transcriptionally and functionally^25,28^. However, our previous studies have not clarified the specific transcriptional changes involved, nor how these changes influence VSMC heterogeneity, cellular states, and define a specific disease-related phenotype concordant with phenotypes observed in human atherosclerotic tissue.

In this study, we leveraged the power of induced pluripotent stem cells (iPSCs) and their vascular derivatives to determine the role of the 9p21.3 risk haplotype in VSMC state transition. iPSCs accurately maintain the genetic background of the donor without carrying the epigenetic marks of the somatic tissue from which they originate, erased during the reprogramming process. Paired with fine genome editing, they provide a unique opportunity to unravel genotype-to-phenotype causal effects at specific genomic regions. Moreover, since they grow in an isolated and controlled environment, they are excellent tools for evaluating cell-autonomous mechanisms in specific cell types based on their genetic repertoire.

Using iPSCs from multiple donors carrying either the risk or non-risk haplotype at 9p21.3, we have unraveled the causal effects of the 9p21.3 CAD risk locus on smooth muscle cell state trajectories. Our analysis provides a gene network map detailing the acquisition of cell state transitions across different genotypes and demonstrates that the transcriptional and functional defects in RR VSMCs are primarily driven by altered expression of *ANRIL*. This study presents several innovative concepts and provides significant progress in understanding the role of the 9p21.3 risk locus in VMSCs.

First, we provide direct evidence that the 9p21.3 locus drives the VSMC state toward an *osteochondrogenic* phenotype. Risk cells maintain SMC identity but show strong upregulation of *SOX9* and *COL2A1*, along with other markers of osteogenic and chondrogenic lineages. Osteochondrogenic SMC states, characterized by loss of contractility and increased expression of *SOX9*, *RUNX2* and *BGLAP*, have been previously reported to be associated with CAD^23^. We previously showed that RR VSMCs have reduced contraction force^25^, and now provide evidence for a significant defect in migration capacity. Importantly, here we show defective migration in iPSC-VSMCs from several donors with risk and non-risk genotypes, as well as in 9p21.3-genotyped primary HCoASMCs, providing strong evidence for a genotype-specific phenotype. Moreover, although RR VSMCs are not enriched in *RUNX2* cells, they show increased calcification propensity, strongly hinting to a role for the 9p21.3 in promoting calcification. Deletion of the 9p21.3 risk haplotype in the same genomic background does not produce a similar signature, providing strong evidence for a genotype-to-phenotype causal effect of the risk allele. Recently, *SOX9* has been linked to vascular aging and the regulation of extracellular matrix composition, leading to arterial stiffness^38^. Moreover, *Lgals3* has been described as a marker of SMC plasticity and linked to a less migratory capacity^34^. In our model, RR VSMCs showed increased expression of *SOX9* and decreased expression of *LGALS3* with concomitant loss of migration. We speculate that the 9p21.3 CAD risk locus alters the vascular wall by reducing VSMC plasticity, ultimately affecting the adaptability of cells to external stimuli and potential damage, increasing vessel sensitivity to recurrent insults and the inability to transition to different phenotypes essential for maintaining healthy vessel function. Moreover, we suggest that the lack of migration in RR VSMCs may contribute to decreased engagement of VSMCs into the fibrous cap, during atherosclerotic plaque formation, promoting plaque instability. Our data are concordant with previous studies linking the 9p21.3 CAD locus to risk of atherosclerotic stroke^58^ and arterial calcification^2^. Leveraging a trajectory analysis, in this study, we define a 9p21.3 risk-specific VSMC state transition and identified numerous transcription factors involved in this transition to investigate further.

Second, using isogenic knockout lines in both risk and non-risk backgrounds at 9p21.3, we provide evidence that VSMCs carrying the risk haplotype at 9p21.3 show a transcriptional signature with minimal overlap with other genotypes and low heterogeneity. Remarkably, the isogenic knockout cells restore a signature similar to non-risk cells, demonstrating that the deletion of the 9p21.3 locus is sufficient to revert their phenotype. We demonstrate the role of 9p21.3 CAD risk locus in driving VSMCs states in a cell-autonomous manner, independent of cues from the surrounding tissue niche. Most studies exploring VSMCs phenotypic modulation have leveraged lineage tracing in mouse models limiting to decipher whether the surrounding tissue or other vascular layers play a role in triggering VSMCs phenotypic switch. Here, we demonstrate that VSMCs carrying the risk haplotype at 9p21.3 have an intrinsic potential to acquire an osteochondrogenic-like state. Hence, we show that the presence of the risk allele is sufficient to guide this cell transition independent of other vascular cells.

Third, in this study, we also identified two new signature genes, *LIMCH1* and *CRABP1*, that, combined with *SOX9* and *COL2A1* expression, define the 9p21.3 risk status. These genes are uniquely expressed in risk VSMCs and are minimally expressed in other genotypes, suggesting that they are controlled by the 9p21.3 risk haplotype. Although *LIMCH1* and *CRABP1* have been previously detected during heart development^59,60^, their roles in VSMC differentiation and state transition have not been described. Importantly, LIMCH1 has been showed to inhibit migration in HeLa cells^61^, supporting a link between high expression of this transcript in RR VSMCs and the concomitant lack of migration. Despite identifying new roles for *LIMCH1* and *CRABP1* in mediating SMC cellular states, further studies are needed to fully characterize their vascular-specific functions. Importantly, we provide evidence that these genes are expressed in *ex vivo* human coronary and carotid arteries, limiting potential artifacts of our system. Importantly, *LIMCH1* and *CRABP1* expression levels were consistently increased in iPSC-VSMCs derived from additional 9p21.3 risk compared to non-risk donors, strengthening the link between the 9p21.3 risk haplotype and upregulation of these signature genes. Using exogenous overexpression of *ANRIL isoform 12*, we previously identified uniquely expressed in risk VSMCs^25^, we demonstrate that *ANRIL* is sufficient to elicit the upregulation of the genes featuring the 9p21.3 risk signature, providing strong evidence that this phenotype is likely induced by the altered expression of ANRIL short isoforms in risk VSMCs.

Our integrative transcriptome analysis, including iPSC-VSMCs and *ex vivo* SMCs, showed that iPSC-derived VSMCs reflect the overall transcriptional profile of *ex vivo* SMCs and recapitulate the heterogeneity of native tissue cells. This highlights the power of this model to explore cellular state dynamics in the vascular wall and exploring mechanisms driving VSMC phenotypic modulation. Interestingly, the RR VSMCs also composed a unique cluster that resembled a continuous and advanced stage of *osteochondrogenic VSMCs* in the pseudotime trajectory. We thus speculate that the atlas may not include the full representations of this extreme phenotype, due to technical challenges in extracting these cells from calcified human plaques or lack of sufficient representation of the risk haplotype at 9p21.3 among the samples. Alternatively, it is possible that our iPSC-based culture system enriches this phenotype to a degree that is not captured *in vivo*. We also cannot exclude the possibility that this extreme phenotype *in vivo* is buffered by other signals within the complex tissue environment.

Our study has some limitations, since it does not provide a temporal projection of the osteochondrogenic signature acquisition during the VSMC differentiation. Additional studies would be required to explore further how the risk VSMCs phenotype is acquired and can be prevented. Moreover, additional validation studies would be required to explore the cellular composition of coronary arteries from human donors carrying the risk haplotype at 9p21.3. Another limitation of this study is that the scRNA-seq dataset is based on iPSC-VSMCs derived from two human donors. However, to address this limitation, we have used iPSC-VSMCs from additional donors and primary human smooth muscle cells to validate our main findings. Expanding the number of lines included in future transcriptomic analyses will further enhance our ability to identify additional 9p21-specific gene signatures.

Overall, our study provides yet undescribed evidence on the role of the 9p21.3 CAD risk locus in altering VSMC state transitions, promoting an osteochondrogenic state. Moreover, we highlight the power of the iPSC-based system to reveal the biology of VSMCs and potentially other disease-relevant cell types to understand the dynamics of cellular trajectories driven by the most impactful CAD genomic locus.

## Non-standard Abbreviations and Acronyms

iPSC: induced Pluripotent Stem Cell
VSMC: Vascular Smooth Muscle Cell
HCoASMC: Human Coronary Artery Smooth Muscle Cell
GWAS: Genome Wide Association Study
CAD: Coronary Artery Disease
lncRNA: long non coding RNA
UMAP: Uniform Manifold Approximation and Projection
GO: Gene Ontology
RR: cells homozygous for the CAD risk haplotype at 9p21.3
RR KO: cells carrying a complete deletion of both risk alleles at the 9p21.3 and isogenic to RR cells
NN: cells homozygous for the non-risk haplotype at 9p21.3
NN KO: cells carrying a complete deletion of both non-risk alleles at the 9p21.3 and isogenic to NN cells.
RR_2: cells homozygous for the CAD risk haplotype at 9p21.3 from a second donor
RR_3: cells homozygous for the CAD risk haplotype at 9p21.3 from a third donor
NN_2: cells homozygous for the non-risk haplotype at 9p21.3 from a second donor
NN_3: cells homozygous for the non-risk haplotype at 9p21.3 from a third donor

## Acknowledgments

The authors would like to thank Dr. Ali Torkamani and Doug Evans at the Scripps Research Translational Institute, Katharine Hubert in Dr. Deneen Wellik’s laboratory and Huan Yang in Dr. Bo Liu’s laboratory at UW-Madison SMPH for initial guidance on sequencing data analysis. The authors utilized the University of Wisconsin – Madison Biotechnology Center Gene Expression Center (Research Resource Identifier – RRID:SCR_017757).

## Supplemental Material

- Expanded Material and Methods

## SUPPLEMENTAL MATERIALS

### Material & Methods

#### Data availability

All data generated via scRNA-seq are deposited in GEO and will be released upon publication and made publicly available. All other data supporting the findings in this study are included in the main article and associated Supplementary Tables.

#### Ethical compliance and cell lines

The authors have complied with all ethical regulations. iPSC lines were previously derived and characterized^25,40^ under a study approved by the Scripps Research Institute IRB (IRB#115676, IRB#07-4714). The cells were transferred and maintained under a study approved by the University of Wisconsin Madison through Material Transfer Agreement (MSN246500). The lines included in this study were obtained from a total of six donors. Lines included in the scRNA sequencing analysis were obtained from two donors described in Lo Sardo *et al*. Cell 2018^25^. iPSC-VSMCs, including unedited clones from additional RR and NN donors (Lo Sardo *et al.*, Nat Biotech 2016^40^), were used to confirm the main findings. HCoASMC were purchased from *Cell Application* and genotyped at the 9p21.3 locus for rs1333049 and rs10757278 with a protocol previously described^25^. A complete list of lines is present in **Table S8**.

#### iPSC derivation, differentiation and maintenance

iPSCs were cultured on Matrigel-coated (Corning 354277) cell culture plates and mTeSR (Stem Cell Technologies cat. 85850) and cultured at 37°C under normoxic conditions (20% O_2_) with 5% CO_2_. iPSCs were differentiated into VSMCs as previously described^25,26^. VSMCs were maintained on VSMC media (DMEM, Gibco 11965092; 10% BenchMark™ Fetal Bovine Serum, Gemini; Glutamax, Gibco 35050061; Penicillin/Streptomucin, Gibco 15140122; MEM Non-Essential Amino Acids, Gibco 11140050). For calcification assay VSMCs were cultured until 80% confluent and fed with Calcification media^39^ for 7 days.

#### ANRIL Isoform overexpression

##### Lentivirus preparation

Reference isoform 12 of ANRIL inserted into a 3rd generation lentiviral vector with tetracycline response element and truncated HIV 3’LTR were generated according to the method described in Lo Sardo et al. 2018. To prepare lentivirus, LentiX HEK293T cells (632180 Clon-tech inc.) (RRID: CVCL_0063) at 60% confluency were transfected with 3rd generation packaging vectors (REV, RRE and pMD 2.G) (RRID: Addgene_12253, Addgene_12251 and Addgene_12259) and TetO_ANRIL 12 or rtTA M2.2 plasmids using the calcium phosphate method. 24 hours after transfection, the media was replaced. Virus was harvested 24 hrs after the media change (48 hrs after transfection). After filtering through 0.45μm low protein binding syringe filters (SLHVM33RS Millipore) supernatant containing the viral particles was stored at -80℃.

##### ANRIL overexpression in iPSC-VSMC Differentiation

iPSC with NNKO background were differentiated to VSMC as previously described, and co-infected with lentiviruses containing rtTA M2.2 and ANRIL 12 or rtTA only, on day 7 of the differentiation. The virus was removed 24 hrs later and the cells were fed with Doxycycline 1μg/mL (Millipore Sigma, Cat. D9891) to induce overexpression until collection on day 17.

#### Single-cell library preparation and sequencing

Isogenic iPSCs (RR, RRKO, NN, NNKO) pairs, for a total of 8 lines, were differentiated in parallel. Terminally differentiated VSMC were collected, and libraries were generated simultaneously, and processed as per 10X genomics single cell preparation instruction. The cells were maintained on ice and subjected to library preparation at the University of Wisconsin–Madison Biotechnology Center’s Gene Expression Center Core Facility (Research Resource Identifier - RRID:SCR_017757) for single-cell RNA library preparation and the DNA Sequencing Facility (RRID:SCR_017759) for sequencing. GEM and library preparation were conducted through the Core using a 10x Genomics Chromium Next GEM Single Cell 3’ Reagent Kit v3.1 and a single-cell G Chip Kit (Lot# 161273). All protocols were followed according to the manufacturer’s instructions. The target cell recovery (TCR) values were determined according to the quality and viability of the samples and calculated for each sample to obtain ∼3000 cells/sample. Sample libraries were balanced for the number of estimated reads per cell and run on an Illumina NovaSeq system on a NovaSeq S1 flow cell at the University of Wisconsin–Madison Biotechnology Center’s Gene Expression Center Core Facility.

Single-cell RNA-seq data were first analyzed by the UW Bioinformatics Resource Center (RRID:SCR_017799) using Cell Ranger pipelines from 10x Genomics (RRID:SCR_023672). The data were demultiplexed with the Cell Ranger mkfastq command wrapped around Illumina’s bcl2fastq. Cell Ranger software (RRID:SCR_017344) was then used to perform alignment, filtering, barcode counting, UMI counting, and gene expression estimation for each sample according to the 10x Genomics documentation. The gene expression estimates from each sample were then aggregated using CellRanger (cellranger aggr) to compare experimental groups with normalized sequencing depth and expression data. This produced a dataset of 27,088 cells.

#### Bioinformatic analysis of single-cell sequencing of iPSC-VSMCs

##### Quality Control

The data were analyzed in R studio (v. 4.2.1) using Seurat V4.3.0 (RRID: SCR_016341). Genes expressed in fewer than 3 cells and cells with fewer than 200 genes expressed were omitted. The data were filtered to remove dying cells, doublets, and empty GEMs using UMI count (5,000-120,000), feature count (2,500-8,500), and percent mitochondrial RNA (<20%). Since the entire dataset was produced in a single batch, with all cells cultured, prepared, and sequenced simultaneously, no batch correction was performed. However, as a QC the Seurat object containing the precomputed linear dimensional reduction PCA was passed to the R package Harmony with default settings to generate a corrected PCA. Harmony components were then employed to perform non-linear dimensional reduction using RunUMAP() and cell clustering by running FindNeighbors() and FindClusters(). This control is presented in the Supplementary material. Pseudobulking was performed to analyze the similarity of different subclones with the same genotype. To pseudobulk, we utilized Seurat::AggregateExpression() to sum up the raw gene counts of all cells from the same sample, resulting in a gene expression profile for each sample. Subsequently, pseudobulk gene counts were subjected to R stats::prcomp() and stats::cor() to perform principal component analysis and Pearson’s or Spearman’s sample correlation analysis, respectively.

##### Normalization, PCA, and clustering

Normalization, principal component analysis (PCA), and clustering were conducted using Seurat V4.3.0. After filtering, counts were normalized via SC transformation with variation from % mitochondrial RNA regressed out, analyzed via PCA, and clustered via the Louvain method. Five principal components and a resolution of 0.4 were used in the final clustering.

##### scRNA data figures

All UMAPS, heatmaps, and violin plots were generated using the Seurat V4.3.0 commands RunUMAP and dimPlot for UMAPs and Doheatmap and Vlnplt for the heatmaps and violin plots, respectively. Dot plots and bar charts were generated with ggplot2 (version 3.4.2) (SCR_014601) DotPlot and ggplot commands.

##### Gene Ontology

Gene Ontology (GO) analysis was conducted using clusterProfiler (version 3.0.4) (RRID:SCR_016884).

##### Trajectory Analysis

Trajectory analysis was conducted using Monocle3 (version 1.3.1) (RRID:SCR_018685). Trajectory heatmaps were generated using the cluster_pseudotime_genes function from the Miller Laboratory GitHub (https://github.com/MillerLab-CPHG/Human_athero_scRNA_meta).

#### Bioinformatic analysis of publicly available datasets

Comparison to published datasets was conducted by downloading the original count matrices and filtering/clustering using the metrics and filters detailed in Table S9. Thresholds for filtering were set to omit excessively high % mitochondrial RNA and unusually high or low UMI and feature counts relative to the bulk of the cells in the dataset to capture the representative majority of cells and omit dead cells, doublets, or empty GEMs. Once the downloaded datasets were clustered and a UMAP was generated, clusters were annotated for cell types using cell type-specific markers provided by the original publications. The resolution for published datasets was empirically estimated to approximate the same number of cell type clusters as in the original publication. The parameters used for analysis are included in **Table S9**.

#### Immunocytochemistry and Alizarin Red staining

Cells were first fixed using 4% paraformaldehyde (15710 Electron Microscopy Science) in PBS. <u>Immunocytochemistry:</u> the fixed samples were incubated with 0.5% Triton X-100 (T8787 Sigma-Aldrich) in PBS for 10 minutes. 5% FBS (100-106 Gemini Bio) in PBS was used for blocking for 1 hour at RT. Primary antibodies against LIMCH1 (NBP2-39006 Novus Biological, RRID: AB_3298404) and/or CRABP1 (sc-166897 Santa Cruz, RRID: AB_10613989) diluted in blocking buffer were incubated for approximately 2 hours at room temperature or overnight at 4°C. After three PBS washes the secondary antibodies Donkey anti-mouse 488 (Alexa-fluor A32766 Invitrogen, RRID: AB_2762823), Donkey anti-rabbit 555 (Alexa-fluor A32794 Invitrogen, RRID:AB_2762834) and DAPI (MBD0015 Sigma‒Aldrich) in blocking buffer were added for 1 hour at RT. Three additional PBS washes were performed. <u>Alizarin Red:</u> Cells were rinsed twice with distilled water and incubated at RT with 2% Alizarin Red S Solution (pH 4.2) (2003999 EMD Millipore) for 30 minutes. Stain was then removed, and cells were washed with distilled water three times. Samples were imaged via fluorescence and light microscopy in a Nikon Eclipse and Alizarin red staining was acquired in a Keyence BZ-X800 (RRID: SCR_023617). Quantification of Alizarin red staining: For each condition (Un and Ca) and cell line, 3–5 images were captured using a Keyence BZ-X800 microscope. The staining intensity was measured using Fiji ImageJ software (RRID:SCR_003070) and normalized to the area covered by the cells. Data are presented as Relative Calcification, calculated as the ratio of the normalized intensity for each cell line to the average signal across all cell lines.

#### RNA extraction, cDNA synthesis, and qPCR

The cell pellets were dissociated using TRIzol (15596018 Invitrogen). RNA was then extracted using a Zymo Direct-zol RNA Miniprep kit (R2052) according to the manufacturer’s protocol. iScript Reverse Transcription Reaction Mix (1708891, Bio-Rad) was used for cDNA synthesis. For qPCR, iTaq Universal SYBR Green Supermix (1725121 Bio-Rad) and SsoAdvanced Universal SYBR Green Supermix (17252721 Bio-Rad) was used on a Bio-Rad CFX OPUS 384 Touch Real-Time PCR machine. GraphPad Prism (RRID: SCR_002798) was used to perform statistical analysis using unpaired t-tests or 1-way ANOVA with Bonferroni correction. A complete list of primers used for qPCR is provided in **Table S10**.

#### Migration assay with Incucyte S3

Migration assays were performed following the scratch wound protocol from Sartorius. ImageLock 96-well plates (BA-04857 Essen Biosciences) were coated with Corning Matrigel hESC-qualified Matrix (354277 Corning) at the recommended concentration of 5µg/well and left to polymerize for 1h at 37C. Cells were cultured in VSMC media and detached using TrypLE Express (12-604-013 Gibco). Cell counts were done using a Countess II automated cell counter (AMQAX1000 Invitrogen) before seeding at 80k/well. Cells were allowed to grow overnight before an Incucyte Woundmaker Tool (4563 Sartorius) was used to create uniform scratch wounds. Cells were washed twice with phosphate buffered saline (14-080-055 Gibco) before 200µg Matrigel was added to each well and allowed to polymerize for 1h at 37C. 100µl media was added to each well after polymerization, and the plate was placed in an Incucyte S3 imaging system (RRID: SCR_023147) for scanning every 2 hours. Images were analyzed with the Incucyte 2020A software package to determine wound density normalized to the initial scratch. GraphPad Prism (RRID: SCR_002798) was used to generate graphs, calculate the area under curve, and perform statistical analysis using unpaired t-tests or 1-way ANOVA with Bonferroni correction.

#### MetaPlaq atlas integration

MetaPlaq data integration, co-embedding and trajectory analyses were conducted as previously described in Mosquera *et al*. 2023^20^. Briefly, we subjected the iPSC-VSMC scRNA-seq datasets to an automated data processing pipeline (scRNAutils) that was also used to process the datasets in MetaPlaq, including doublet and ambient RNA removal, filtering for high quality cells, and normalization to account for variability in sequencing depth across cells. PCA was used for dimensionality reduction of the normalized counts matrix and Shared-Nearest-Neighbors (SNN and Louvain community detection was used for clustering in Seurat v4. Clustering results were visualized using UMAP embeddings. Seurat objects were batch integrated using reciprocal PCA, which was shown to optimize biological preservation and removal of unwanted technical batch effects. Clusters were annotated using the original labels in MetaPlaq^20^.

#### Statistical analysis

Statistical analyses are detailed in the corresponding Figure Legends showing number of samples and number of independent experiments. Data were analyzed using unpaired t-test or one-way ANOVA with Bonferroni multiple comparison post hoc test as specified in the corresponding figure legends. Differential expression analysis relative to scRNAseq dataset were analyzed via non-parametric Wilcoxon rank sum test.

---

**Figure S1.**
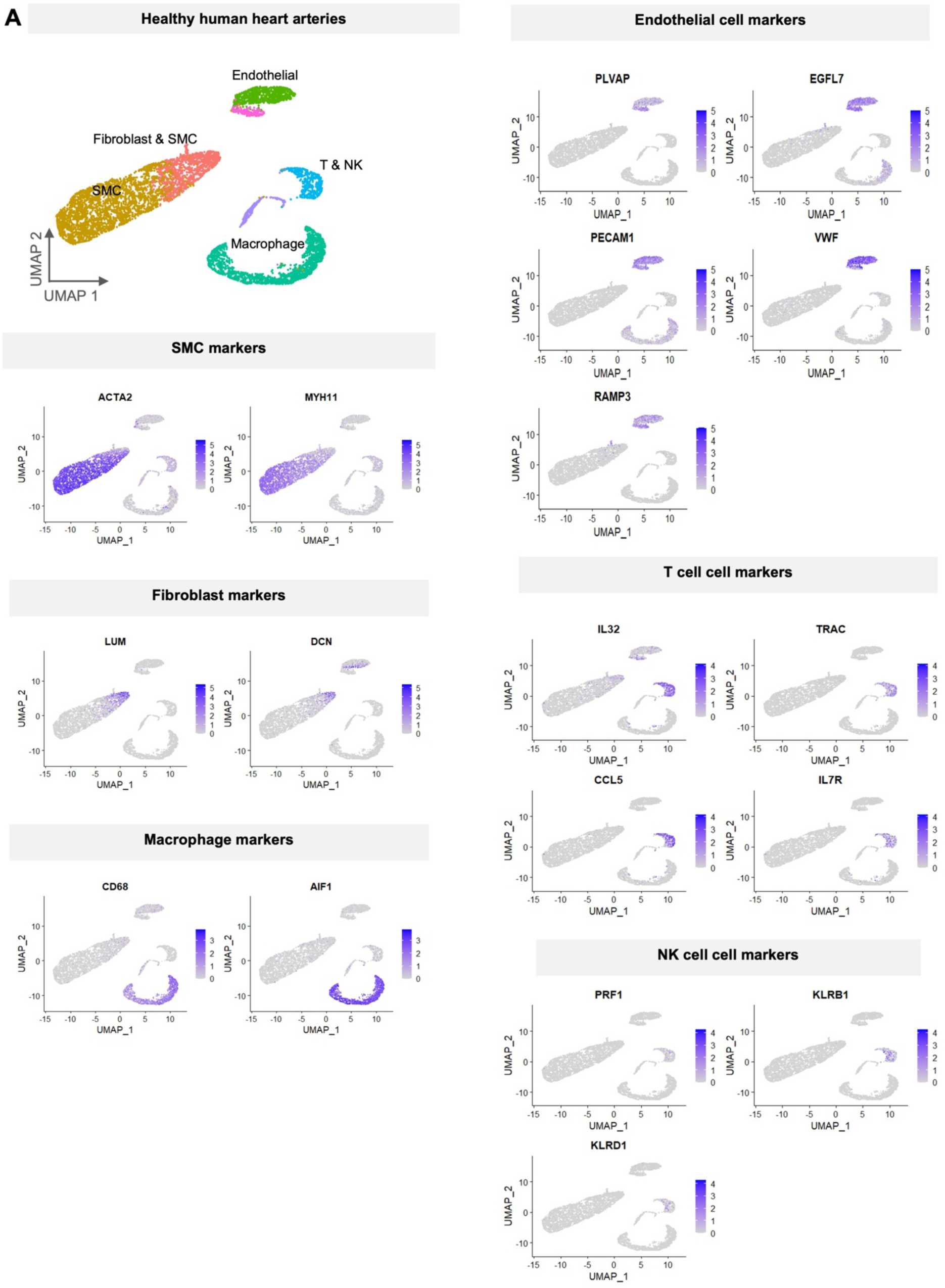
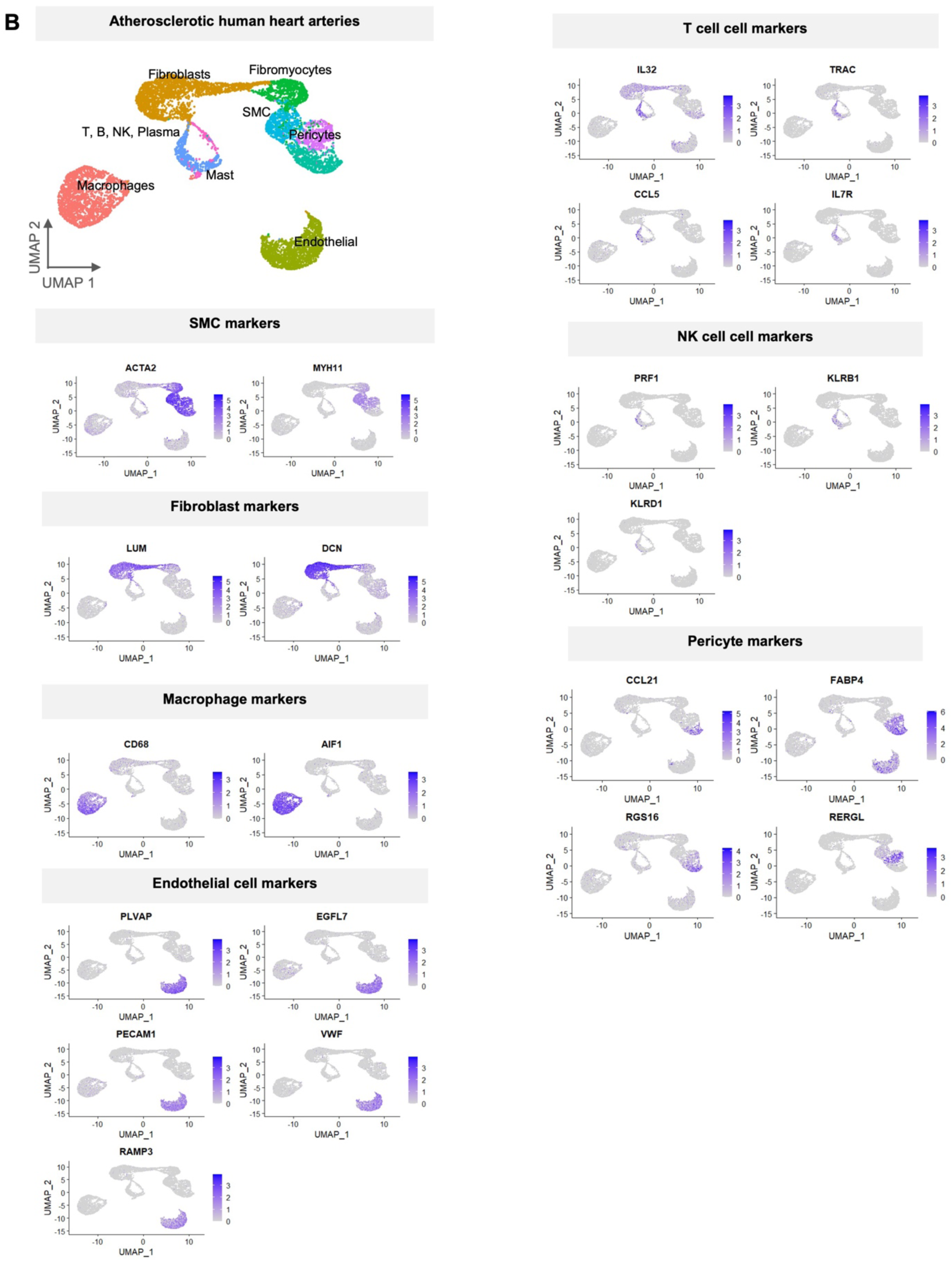
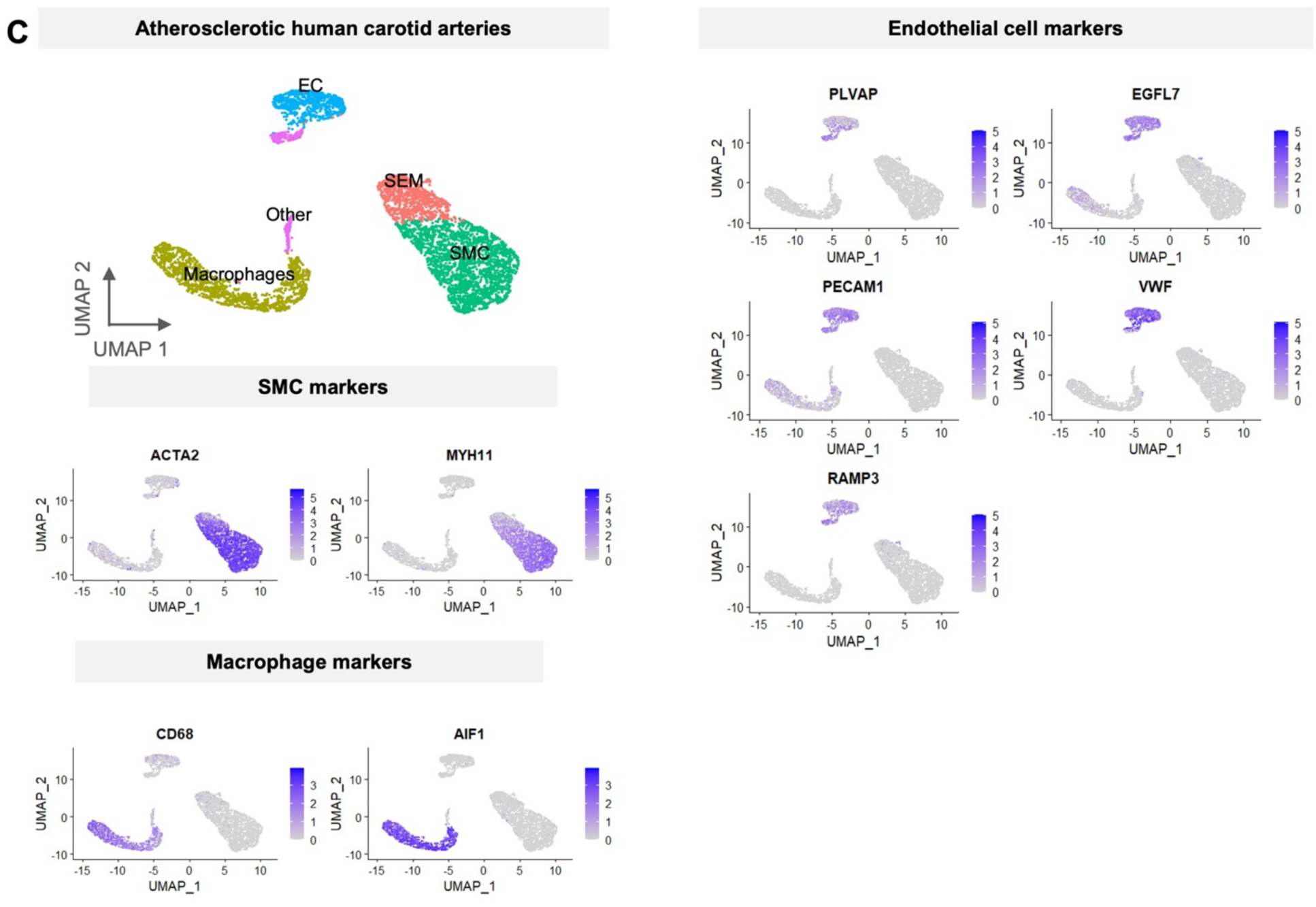
Analysis of publicly available scRNA sequencing dataset from human arteries. UMAPs showing cell type annotations and cell type-specific markers for healthy human artery data from Hu et al., 2021 **(A);** Wirka et al., 2019 **(B)** and Pan et al., 2020 **(C)**.

**Figure S2.**
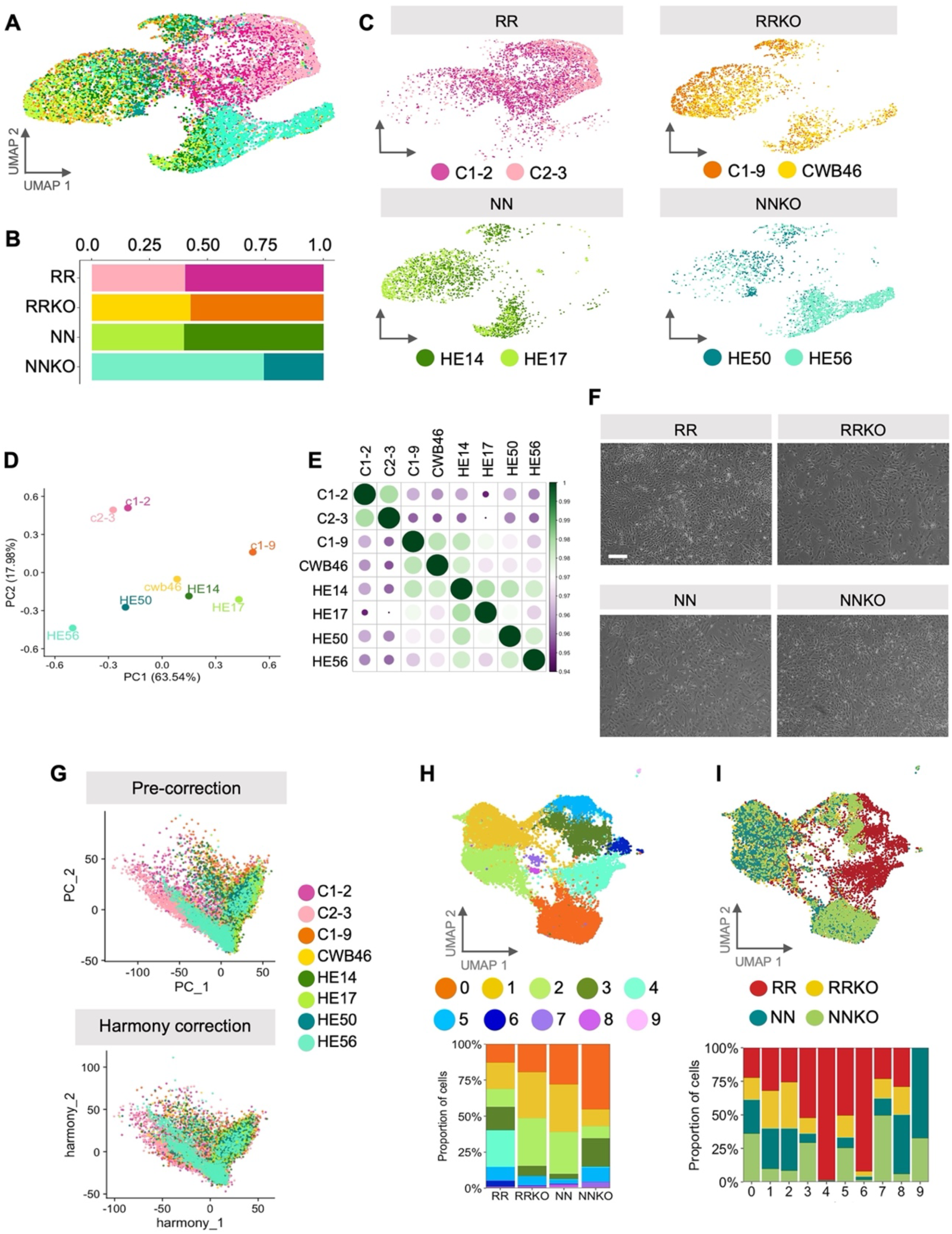
iPSC-VSMCs data QC. **(A)** UMAP of the iPSC-VSMCs scRNA sequencing dataset colored based on individual cell lines (two per genotype). **(B)** Bar chart showing the contribution of each clone to the four genotypes (RR, RRKO, NN, NNKO). **(C)** UMAPs of iPSC-VSMCs divided by genotype and colored by clone contribution. **(D)** Principal Component Analysis (PCA) of the scRNA sequencing dataset after pseudobulk. **(E)** Spearman correlation calculated for all lines included in the scRNAseq. **(F)** Representative photographs of cells of each genotype at D17. Scale bar is 200 μm. **(G-H-I)** PCA and clustering of data after Harmony correction.

**Figure S3.**
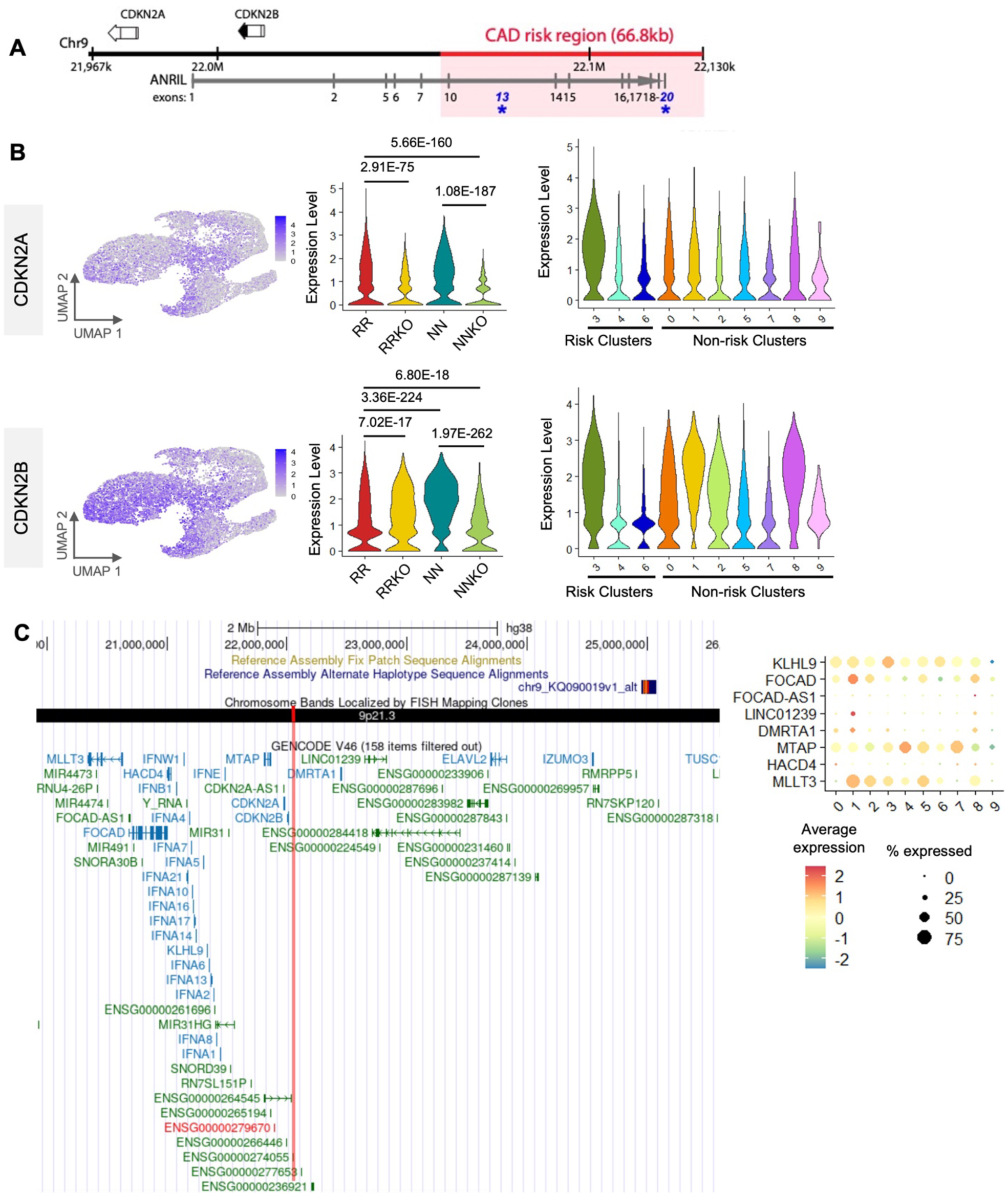
Expression of genes proximal to the 9p21.3 CAD risk locus. **(A**) Diagram of 9p21.3 CAD risk locus with nearest protein-coding genes *CDKN2A/B* and overlapping lncRNA *ANRIL* marked. **(B)** UMAPs of iPSC-derived VSMC single cell data show the expression *CDKN2A* and *CDKN2B* (left), and violin plots show quantification of the same genes across genotypes (middle) and clusters (right). **(C)** UCSC browser view of the 9p21.3 region with all genes annotated and the CAD risk region marked by the red line (left). Bubble plot of the genes from the region (within ∼2 Mb upstream and ∼3.2 Mb downstream) that were detected in the single-cell data (right).

**Figure S4.**
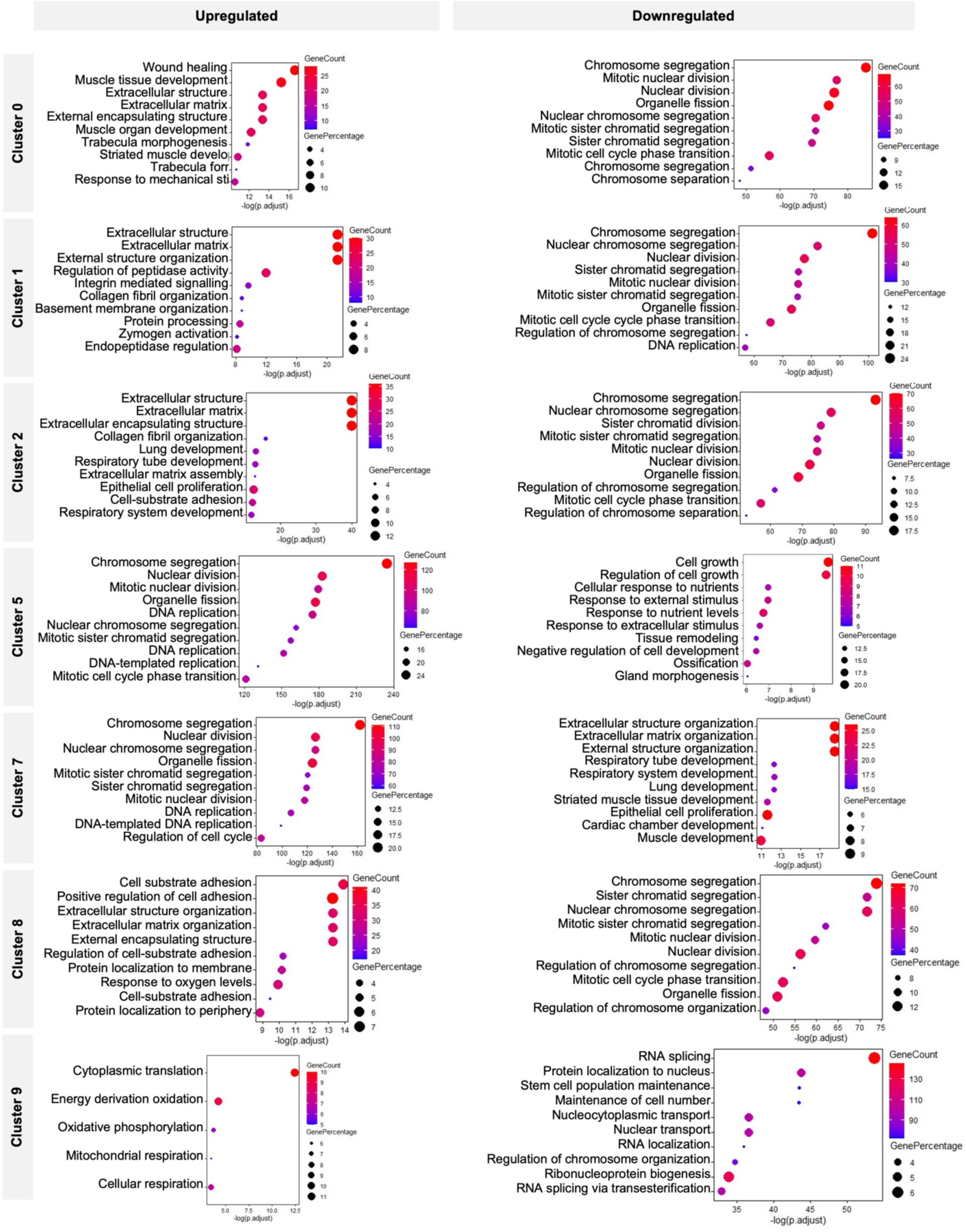
Gene Ontology enrichment analysis for non-RR clusters. Left hand column shows terms enriched in upregulated genes, right shows terms enriched in downregulated genes. Gene ontology analysis conducted with clusterProfiler.

**Figure S5.**
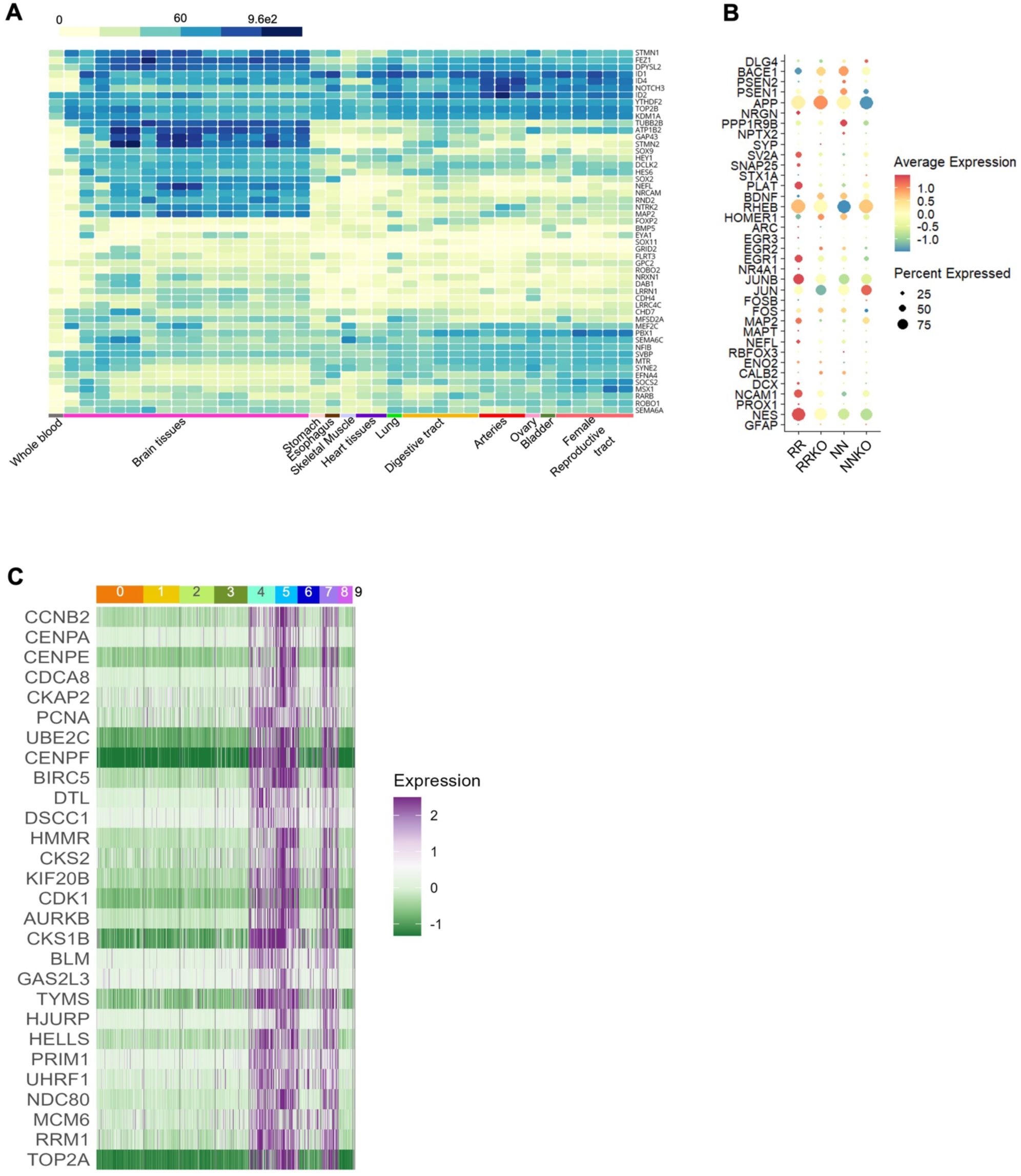
Gene expression profile of axonogenesis and neural projection-related genes and cell cycle-related genes. **(A)** Heatmap of expression of genes associated with neuronal-related gene ontology terms across brain and VSMC-rich human tissues (Generated with the Genotype-Tissue Expression (GTEx) Portal) (Lonsdale et al. 2013). “Female reproductive tract” category includes vagina, uterus, fallopian tube, endocervix, ectocervix. **(B)** Bubble plot showing expression of known neuronal markers across genotypes. **(C)** Heatmap showing the expression of cell cycle related genes across all clusters.

**Figure S6.**
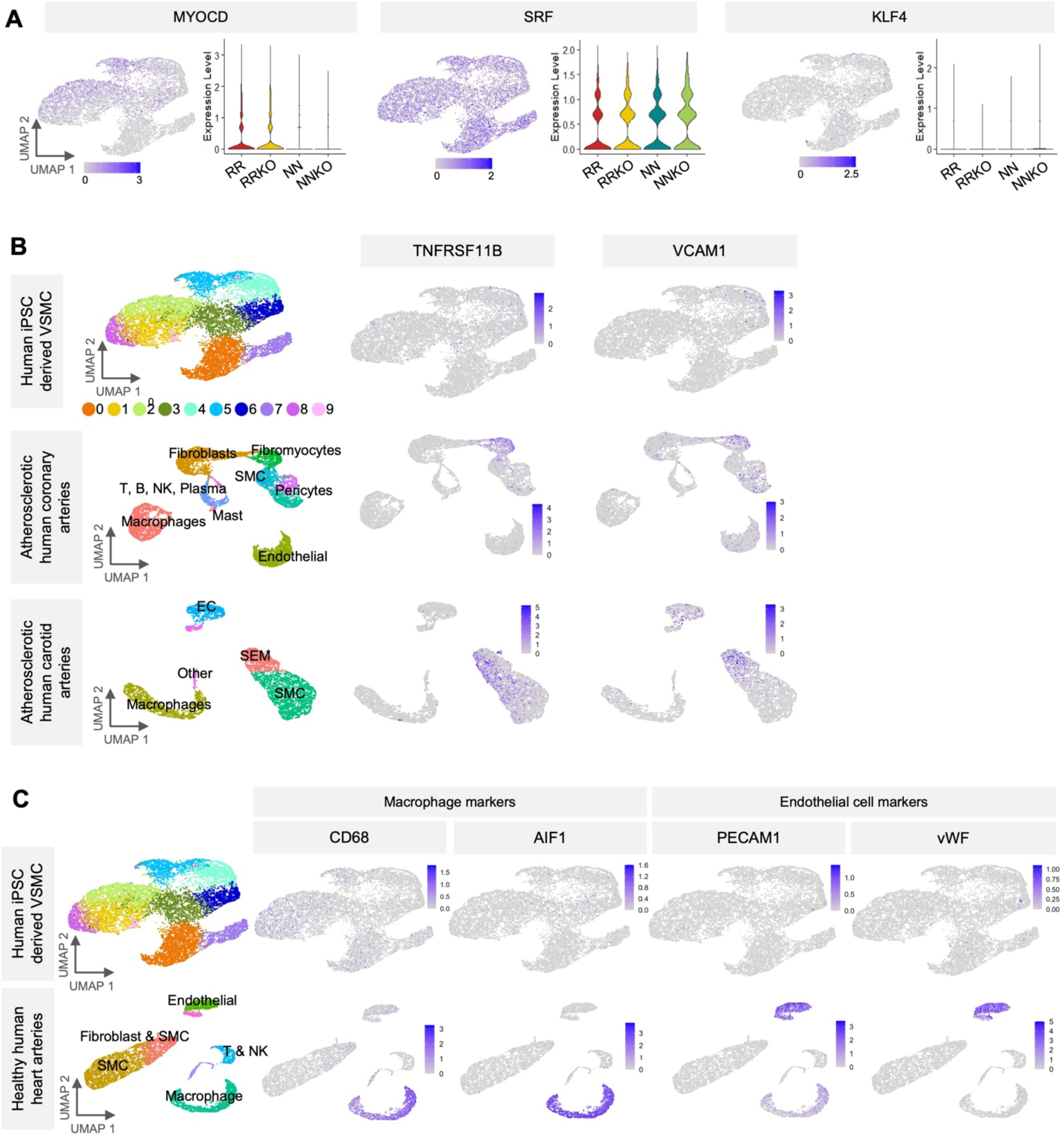
Analysis of known modulators of VSMCs phenotypic switch and known CAD-related VSMCs states. **(A)** UMAPS and violin plots showing expression for three VSMC phenotype modulators, *MYOCD*, *SRF*, and *KLF4*. **(B)** iPSC derived VSMC (top row) compared to atherosclerotic coronary arteries (Wirka at el. 2019, center row) and atherosclerotic carotid arteries (Pan et al. 2020, bottom row). Left hand UMAPs colored by clusters and labeled with cell types, other UMAPs colored according to expression of the gene named above the UMAP. **(C)** Comparison of iPSC-VSMC (top row) to endogenous healthy artery data (bottom row). Two examples of gene markers given for macrophages and endothelial cells.

**Figure S7.**
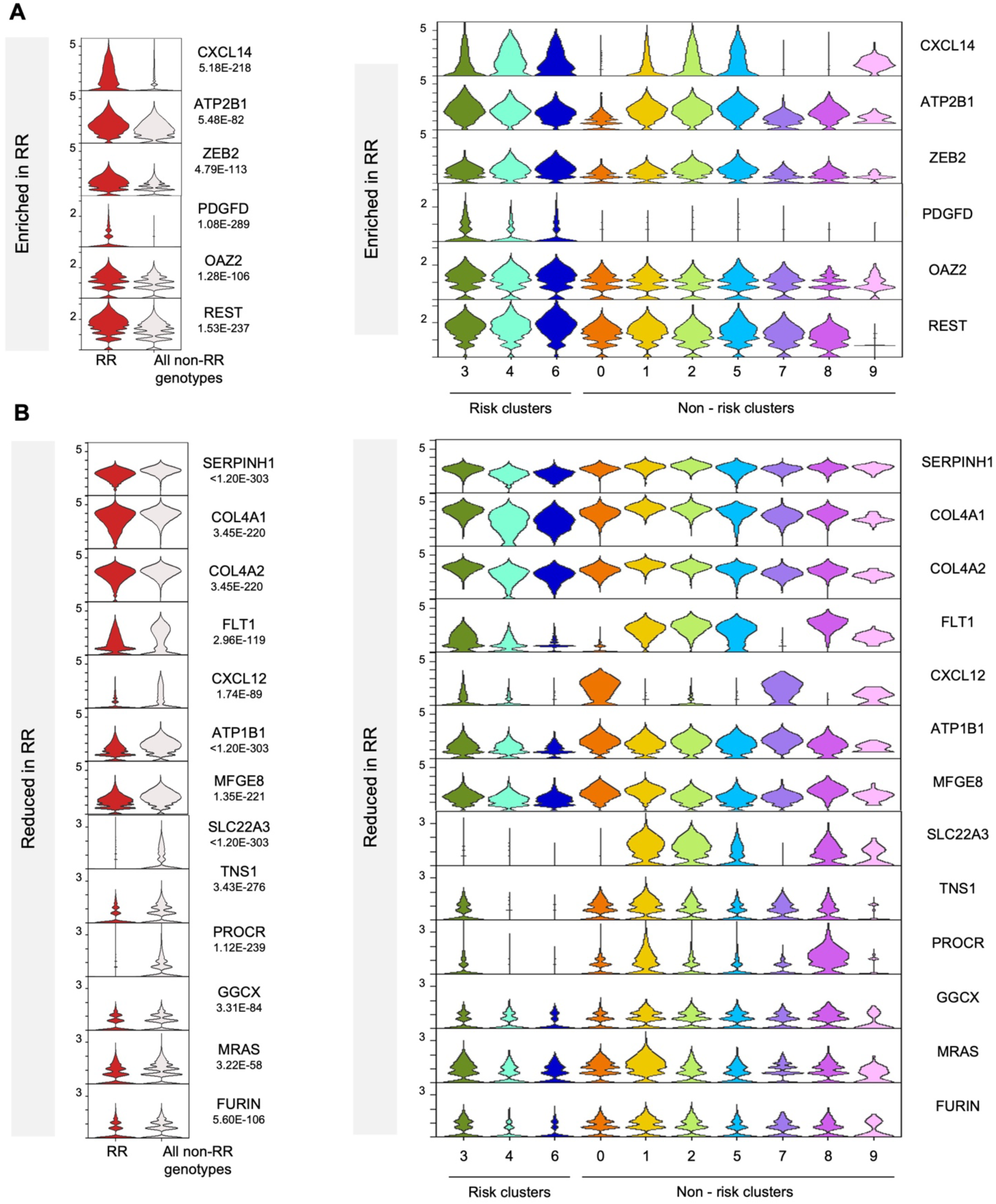
Analysis of a set of CAD-related genes. **(A-B)** Violin plots showing the expression of known CAD genes in RR vs all other genotypes with *p*-values calculated with non-parametric Wilcoxon rank sum test (left). Violin plots of the same genes, showing expression across individual clusters (right). Genes enriched in RR are shown in **A** and genes reduced in expression in RR are shown in **B**.

**Figure S8.**
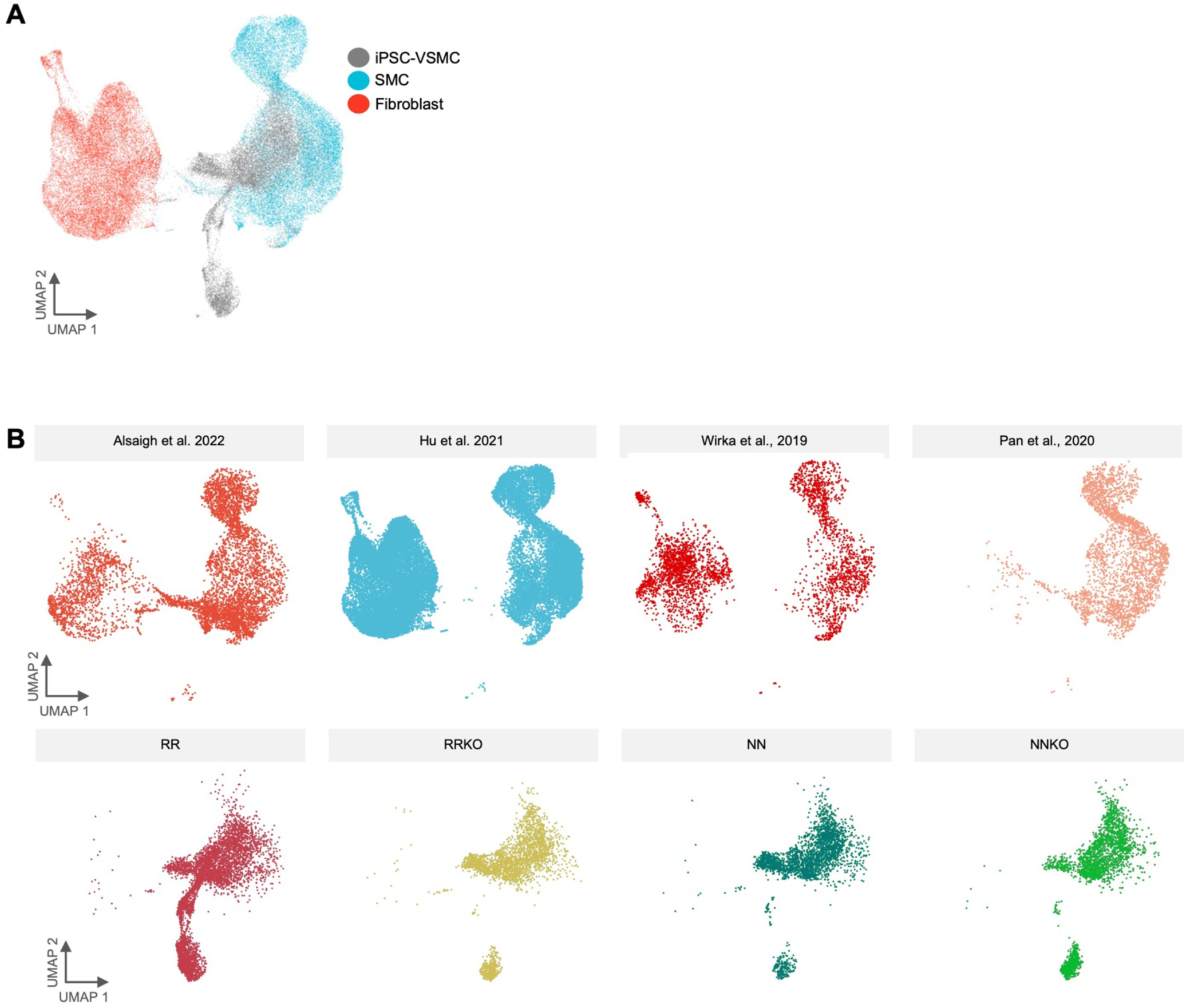
Breakdown of harmonized integration of iPSC-VSMC and SMC and Fibroblasts from the Metaplaq atlas. **(A)** Integration of iPSC-VSMC with Metaplaq atlas Fibroblasts and SMC, colored by cell type. **(B)** Breakdown of harmonization with iPSC-VSMC and Metaplaq atlas SMC/Fibroblasts by contributing dataset (top row) and by iPSC-derived VSMC genotype (bottom row).

**Figure S9.**
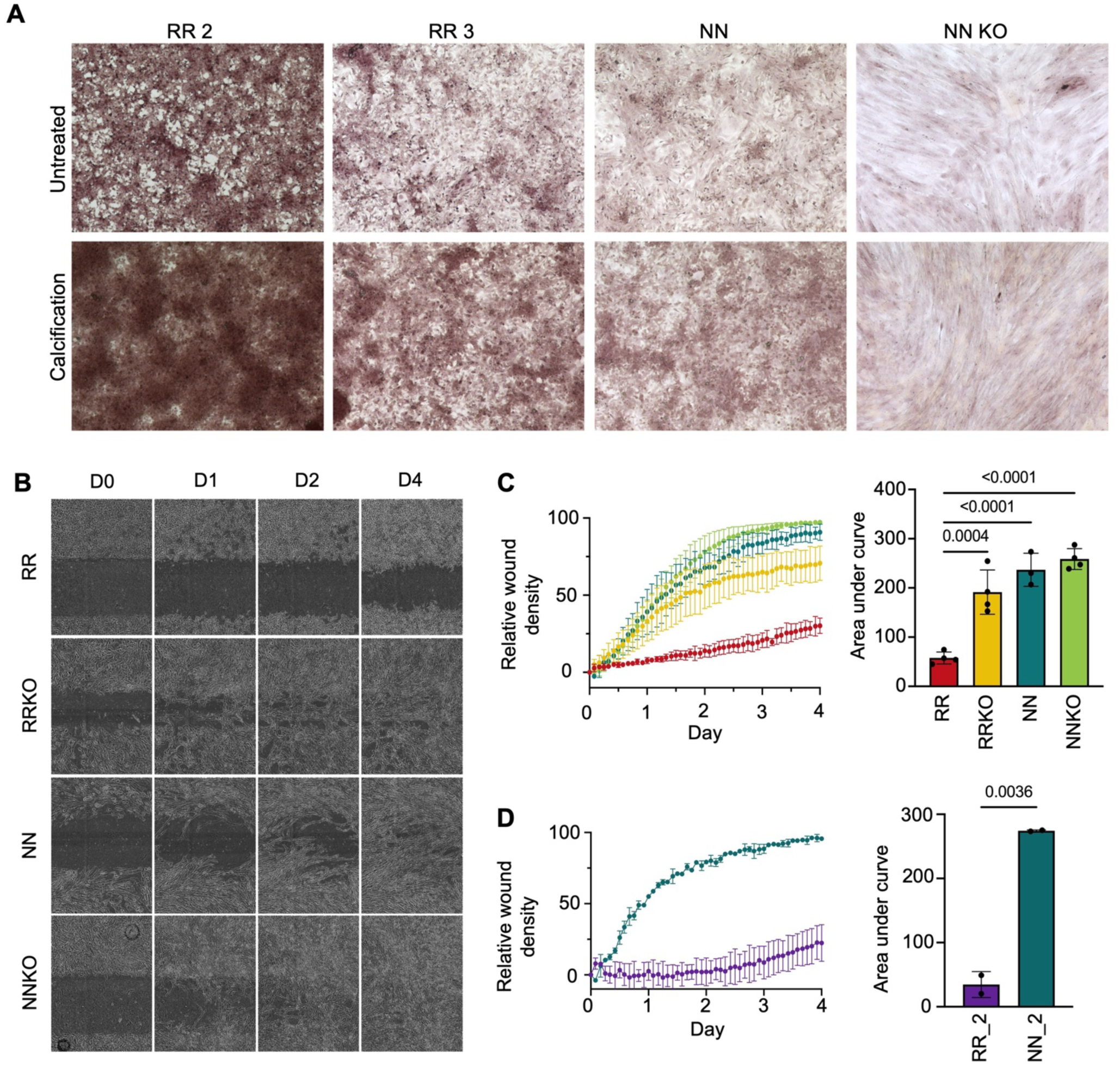
Calcification and migration assays. **(A)** Representative pictures of Alizarin red staining for iPSC-VSMC derived from additional RR donors (RR_2, RR_3) and NN and NNKO lines presented in Figure 4G right panels. **(B)** Photographs of iPSC-VSMCs migration through a scratch assay over time, related to Figure 4H-I. **(C-D)** Breakout of the graph presented in Figure 4I, showing the data from the isogenic RR RRKO and NNKO; N=4 each genotype **(C)** and the additional RR_2 and NN_2 donors (N=2 each genotype) **(D).** Migration assay is plotted over time (left) and area under the curve (AUC) quantification of the assay (right). *P*-value calculated via one-way ANOVA with Bonferroni correction in C, with each sample compared to RR. Unpaired t-test in D.

**Figure S10.**
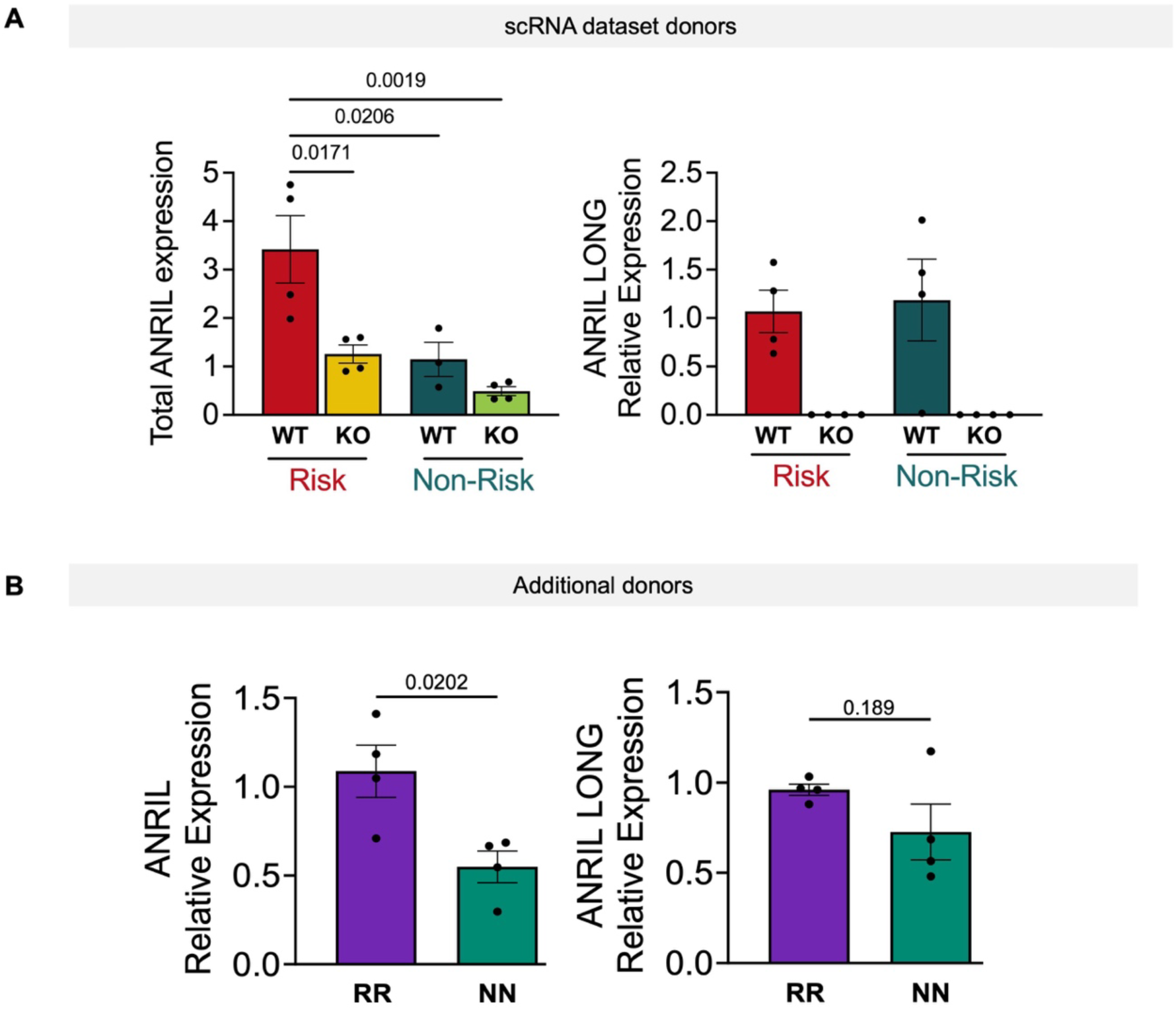
ANRIL isoforms enrichment in RR cells. **(A)** qPCR quantification of total ANRIL (left) and ANRIL long isoforms (right) in cells used to generate the single-cell dataset presented in this work. *P*-values calculated via One-way ANOVA with Bonferroni correction. **(B)** qPCR quantification of total ANRIL (left) and ANRIL long isoforms (right) in iPSC-VSMCs from two additional RR donors (RR_2, RR_3) and two additional NN donors (NN_2, NN_3). *P* values calculated via Unpaired t-test.

